# Multi-modal ion-transport function of potassium channelrhodopsins

**DOI:** 10.64898/2026.09.30.755824

**Authors:** Ryotaro Shimamura, Moe Kikuchi, Toshiki Nakamura, Shuma Tajima, Kazuhiro J. Fujimoto, Masahiro Sugiura, Jinichiro Tabe, Keito Minamisawa, Rei Abe-Yoshizumi, Tatsuro Nishikino, Shoko Hososhima, Takeshi Yanai, Yuji Furutani, Satoshi P. Tsunoda, Hideki Kandori

**Author notes:** Corresponding authors: Satoshi Tsunoda and Hideki Kandori.

## Abstract

Light-gated K^+^ channelrhodopsins (KCRs) are recognized as an ideal neuronal silencing tool in optogenetics. KCRs contain functionally important residues for a light-driven H^+^ pump bacteriorhodopsin, but their roles have been unclear. Utilizing electrophysiological methods, we demonstrate that KCRs function as a light-driven H^+^ pump in the absence of K^+^. Transport experiments using pH electrodes further elucidated the function of the H^+^-pump, which operates even in the presence of K^+^. The hypothesis that the H^+^ pump exerts a blocking effect on K^+^ transport in KCRs has been postulated. Furthermore, we ascertained that there was conversion of KCRs from the H^+^ pump into the H^+^ channel by light, yet this conversion was never reversed. However, the addition of K^+^ resulted in the formation of the K^+^ channel. KCRs exhibit three distinct functional states, H^+^ pump, H^+^ channel, and K^+^ channel. These multifaceted functions are modulated by environmental ions and light.

---

The advent of optogenetics can be traced back to the discovery of channelrhodopsins^1,2^, which elucidated that the application of light triggers neuronal excitation through cellular depolarization, and neuronal silencing through cellular hyperpolarization^3,4^. Microbial rhodopsins have played a pivotal role in optogenetics^5,6^. In this area of study, light-gated cation channels^7–11^ and light-driven inward proton pumps^12,13^ have been utilized as optogenetic tools, leading to cellular depolarization. Conversely, cellular hyperpolarization is achieved through the actions of light-driven chloride pumps^14^, proton pumps^15^, and sodium pumps^16^, or by light-gated anion channels^17^ and potassium channels^18^ (Fig. 1a). Among the various hyperpolarization tools, light-gated potassium channelrhodopsins (KCRs) have been the most anticipated due to the fact that neuronal excitation and silencing accompany the transports of Na^+^ and K^+^, respectively, under physiological conditions^19–23^. However, their practical application faces major biophysical limitations. Under continuous or high-intensity illumination, many KCR variants exhibit a dynamic decline in their K^+^-to-Na^+^ (P_K_/P_Na_) permeability ratio^23^. This ion-selectivity shift allows a transient Na^+^ influx that counteracts hyperpolarization, triggering undesirable depolarization and action potentials-a phenomenon known as the inhibition-to-activation transition.

**Fig. 1:**
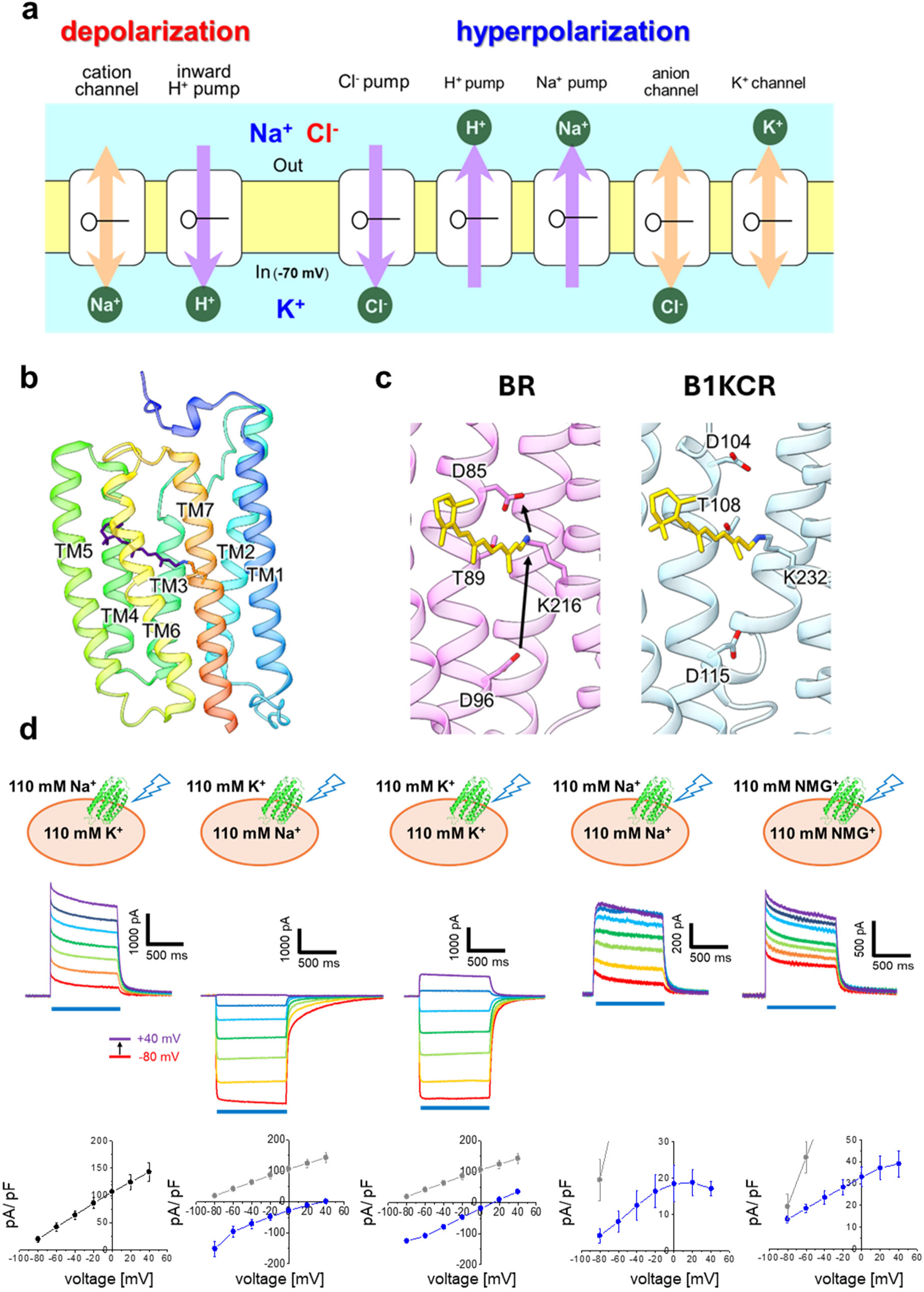
Ion-transporting rhodopsins and a potassium channelrhodopsin, B1KCR. (**a**) Ion-transporting microbial rhodopsins as optogenetic tools. Neuronal excitation occurs by cellular depolarization, for which light-gated cation channels and light-driven inward proton pumps are used in optogenetics. Neuronal silencing is driven by cellular hyperpolarization, for which light-driven chloride pumps, proton pumps, and sodium pumps, or by light-gated anion channels and potassium channels are used in optogenetics. (**b**) The ribbon model of B1KCR is shown in a rainbow color scheme (PDB code: 8JXO), and the all-*trans* retinal is colored in indigo. (**c**) Proton transport pathways associated with the third transmembrane motif and retinal in BR (PDB code: 1M0M) and B1KCR. (**d**) Electrophysiological characterization of B1KCR under different ionic conditions. Whole-cell patch-clamp recordings were performed in B1KCR-expressing ND7/23 cells. The intracellular and extracellular ionic compositions are schematically illustrated. Representative photocurrent traces recorded under five ionic conditions are shown. Blue bars indicate light illumination (480 nm, 6.4 mW mm^-2^). The membrane potential was stepped from −80 to +40 mV in 20-mV increments. Corresponding current–voltage (I–V) relationships are shown.

The transport of ions by microbial rhodopsins occurs in either a passive or active manner. It is important to note that there are substantial differences between channels and pumps^24–26^. General ion channels are defined as pore-forming membrane proteins, whose open/close state is governed by a gate mechanism (Extended Data Fig. 1a)^24,26^. In contrast, the operation of general ion pumps entails the vectorial transport of ions, a process that is facilitated by the application of energy. The underlying mechanism of this process can be elucidated through the framework of the alternating access model (Extended Data Fig. 1b)^25–27^. Consequently, alterations in protein structural for functional processes are more profound in pumps than in channels. In the case of rhodopsins, structures observed in the dark bear a resemblance to channel and pump rhodopsins (Fig. 1b). The presence of multiple gates serves to distinguish between channel and pump functions (Extended Data Fig. 1, c and d).

These rhodopsins must have been optimized for each function during the course of evolution, as is the case for many other proteins. This tendency leads to the assumption that each rhodopsin processes a distinct function^5,6,28–30^. In this report, we revealed the multi-modal ion-transport function of KCRs. It has been established that functionally significant residues in a light-driven H^+^-pump bacteriorhodopsin (BR), characterized by a DTD motif (D85, T89, and D96), exhibit high levels of conservation in cryptophyte channelrhodopsins^31–33^, including KCRs^17,34–40^. In BR, D85 and D96 act as a H^+^ acceptor and donor, respectively, during uni-directional H^+^ pump (Fig. 1c and Extended Data Fig. 1e)^5,6,41–45^. Conversely, the role of the DTD motif in KCRs was not clearly defined. The present electrophysiological study revealed that (i) KCRs become a light-driven H^+^ pump in the absence of K^+^, in which the DTD motif plays an important role as is the case in BR. Subsequently, transport experiments using pH electrodes revealed that (ii) the H^+^-pump function of KCRs occurs even in the presence of K^+^, thereby suggesting that the H^+^ pump is responsible for the termination of K^+^ transport in KCRs. We further revealed (iii) the transition from H^+^ pump into H^+^ channel in the absence of K^+^, which is analogous to a bleaching process in our visual rhodopsins. These findings offer novel insights into ion-transporting rhodopsins. A more profound understanding of multi-modality in KCRs will facilitate the development of new optogenetic tools.

## Light-gated K^+^ channelrhodopsins are light-driven H^+^ pumps in the absence of potassium ions

In this study, we selected a KCR from *Bilabrum sp.* (B1KCR) for electrophysiological measurements^35^. The structure of B1KCR is reported in the dark^40^, which is analogous to the structures of other KCRs and BR (Extended Data Fig. 2a). The retinal Schiff base region of BR contains two positive and negative charges, while R82 in BR is replaced with tryptophan in KCRs, constituting the K^+^ selectivity filter (Extended Data Fig. 2b). As B1KCR was reported to aggregate substantially in intracellular compartments when expressed in mammalian cells^35^, we prepared truncated proteins at the C-terminal side as well as the LR-tagged proteins (Extended Data Fig. 2c). Consequently, the truncated protein at position 292 exhibited enhanced membrane targeting and substantial photocurrents (Extended Data Fig. 2d and e).

We then proceeded to apply whole-cell patch-clamp recordings to B1KCR-expressing ND7/23 cells by modifying the intracellular and extracellular ionic compositions. Photocurrents were observed in both outward and inward directions for K^+^_in_/Na^+^_out_ and Na^+^_in_/K^+^_out_, respectively, indicative of KCR (Fig. 1d and Extended Data Table 1). Based on the Goldmann- Hodgkin-Katz voltage equation, P_K+/Na+_ was determined to be 43.4 ± 5.8 in B1KCR, which falls between the range of HcKCR1 (18 ± 2) and WiChR (80 ± 40)^35^. In the case of K^+^_in_/K^+^_out_, the voltage-dependent photocurrent exhibited asymmetries, which was also the case for other KCRs^35^. We subsequently conducted an experiment with the case of Na^+^_in_/Na^+^_out_, and observed exclusively positive photocurrents under the tested membrane voltage. This phenomenon was more clearly discernible when metal cations were fully removed (NMG^+^_in_/NMG^+^_out_) (Fig. 1d), suggesting that B1KCR facilitates the outward transport of H^+^.

Electrophysiological study demonstrated that B1KCR facilitates the K^+^ transports, yet in the absence of K^+^, it functions as a light-driven H^+^ pump (Fig. 2a). Subsequently, we proceeded to assess the functionality of other KCRs and channelrhodopsins (Extended Data Fig. 3a), in which the DTD motif exhibited full conservation in KCRs and cryptophyte cation channelrhodopsins (Fig. 2b and Extended Data Fig. 3b). Conversely, as *Chlamydomonas reinhardtii* channelrhodopsin 2 (CrChR2) and *Guillardia theta* anion channelrhodopsin 1 (GtACR1) contain the ETH and STL motif, respectively, the DTD motif was introduced into CrChR2 and GtACR1. Photocurrents were recorded under physiological (K^+^_in_/Na^+^_out_) and NMG^+^ /NMG^+^ conditions. Three KCRs, HcKCR1, HcKCR2, and WIChR1, exhibited H^+^- pump activity similar to that of B1KCR (Fig. 2c and Extended Data Table 1). Conversely, in three cation channelrhodopsins (HcCCR, GtCCR4, and ChRmine), H^+^-pump activity was found to be less pronounced in comparison to KCRs. In fact, an inward photocurrent was recorded at -80 mV of HcCCR, which is more indicative of H^+^ channel. In the case of CrChR2, the H^+^ pump signal was found to be minimal. The abolition of photocurrents in CrChR2-DTD and GtACR1-DTD suggests the importance of their respective motif’s function within each rhodopsin.

**Fig. 2:**
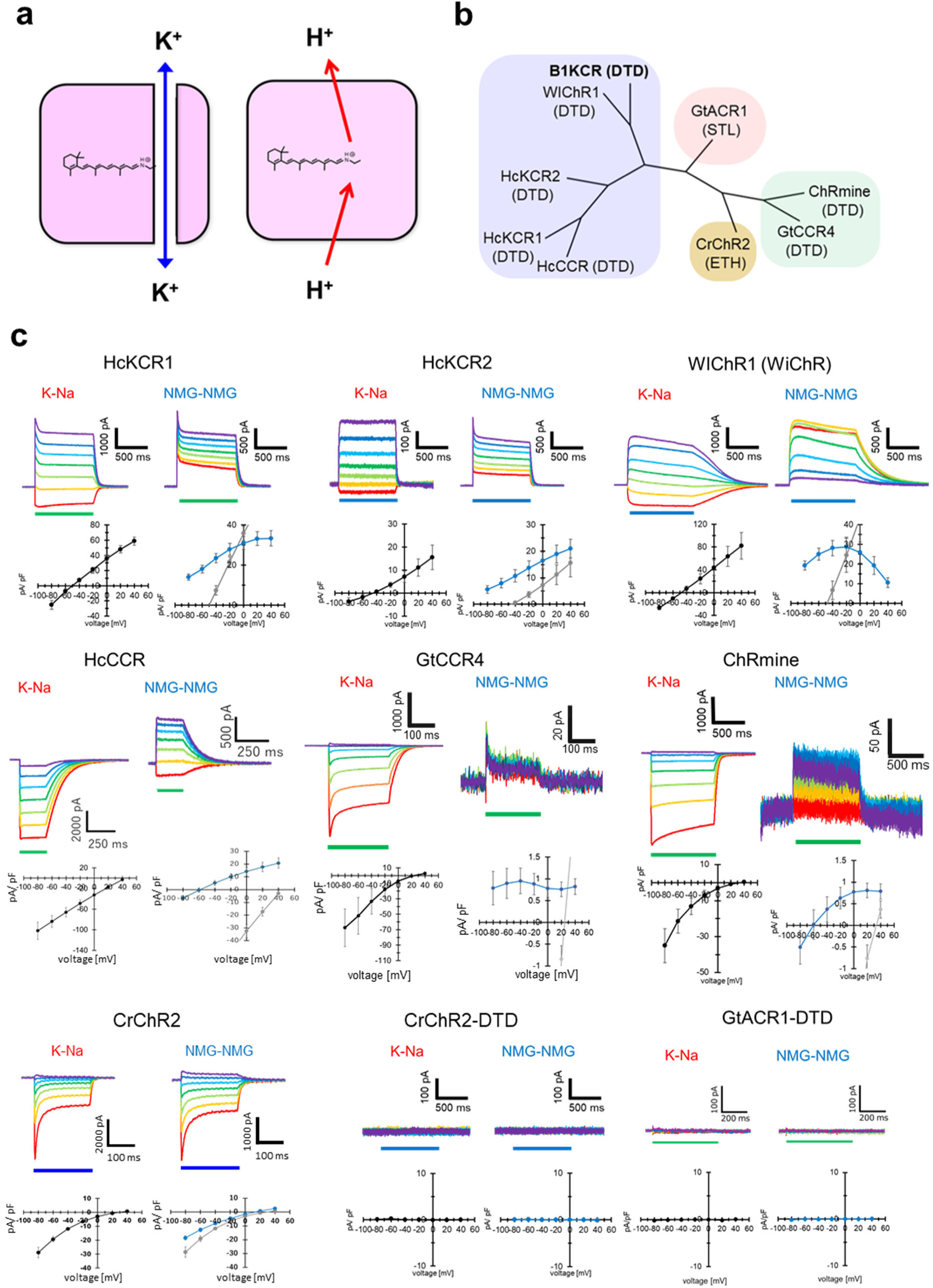
Potassium channelrhodopsins are light-driven proton pumps in the absence of potassium ions. (**a**) Summary of ion-transport in B1KCR. B1KCR is a light-gated potassium channelrhodopsin in the presence of K^+^. In the absence of K^+^, B1KCR becomes a light-driven proton pump. (**b**) Phylogenetic tree of the channel rhodopsins. The TM3 motif is shown in parentheses. (**c**) Photocurrents of channelrhodopsins in the presence and absence of metal cations. Representative photocurrent traces of selected channelrhodopsins and their mutants are shown. K–Na: Recordings obtained with 110 mM K^+^ in the intracellular solution and 110 mM Na^+^ in the extracellular solution. NMG–NMG denotes recordings obtained with 110 mM NMG^+^ in both the intracellular and extracellular solutions. Blue or green bars indicate light illumination at 480 nm (6.4 mW mm^-2^) or 530 nm (1.9 mW mm^-2^), respectively. The membrane potential was stepped from −80 to +40 mV in 20-mV increments. Corresponding I–V relationships are shown. Black or gray indicates recordings obtained in the presence of metal cations, whereas blue indicates recordings obtained in their absence.

It should be noted that the H^+^-pump measurements were conducted in the absence of a pH gradient (pH 7.2_in_/pH 7.2_out_). The H^+^ pump drives vectorial transport of H^+^ against the concentration gradient^25,27,46,47^. Therefore, the same measurements were tested by lowering extracellular pH at 6.0. As a result, positive photocurrents were exclusively obtained for B1KCR, HcKCR1, HcKCR2, and WIChR1 (Extended Data Fig. 4a). In order to quantitatively analyze the properties of the pump, we define the driving force as the membrane potential at which the current reaches zero. The driving forces were analogous between pH 7.2_out_ and pH 6.0_out_ for all KCRs (Extended Data Fig. S4b). Consequently, all KCRs facilitate the outward movement of H^+^ in the absence of K^+^.

There have been some reports on the H^+^-pump function in channelrhodopsins. Photocurrents were observed from CrChR2 in the absence of any ion gradient and electrical potential difference when expressed in electrofused giant HEK293 cells or reconstituted on planar lipid membranes, which was interpreted as an outwardly driven H^+^ pump^48^. In the first paper of KCR, patch clamp recordings of HcKCR1 under NMG^+^ /NMG^+^ conditions yielded photocurrents analogous to those observed in the present study^18^, and outwardly directed active H^+^ transfer has been discussed^39^. Notwithstanding the aforementioned reports, a considerable number of prior experiments have been conducted in the presence of metal cations.

## Ion-transport mechanism in potassium channelrhodopsins

KCRs are a bifunctional ion transporter that is dependent on K^+^. In the presence and absence of K^+^, KCRs transport K^+^ and H^+^, respectively, in passive and active manners. As channels and pumps are distinct molecular machineries, the molecular mechanisms of each transport process are a subject of intrigue. The mechanism of selective transport of K^+^ has been the subject of extensive study for KCRs^17,34–40,49–52^; however, this is not the case for B1KCR which absorbs ble light (Extended Data Fig. 4c). We thus proceeded to mutate B1KCR, which is comprised of three regions; the extracellular K^+^ filter region, the Schiff base region, and the cytoplasmic region (Fig. 3a).

**Fig. 3:**
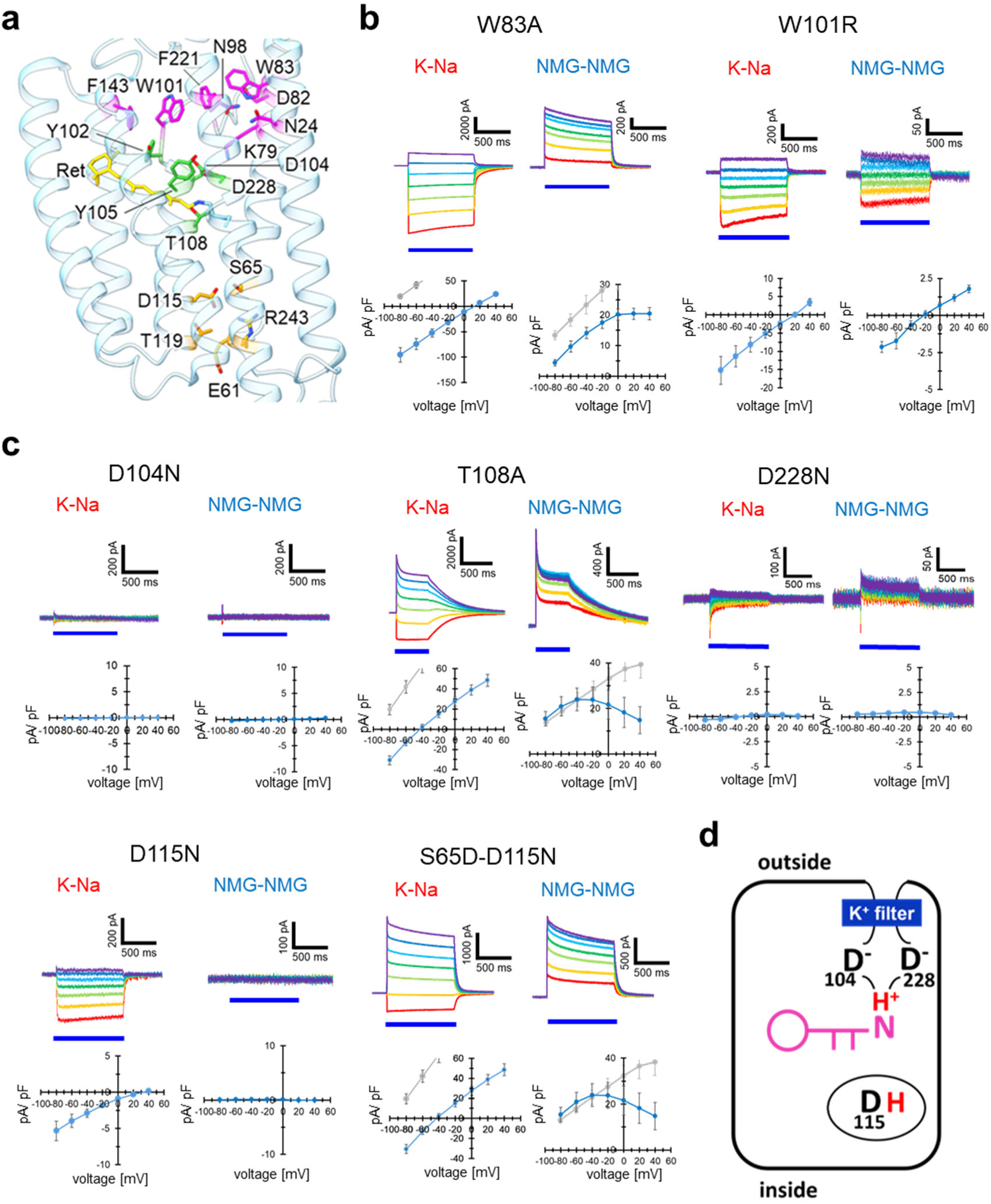
Ion-transport properties of B1KCR studied by use of mutants. (**a**) Amino acid positions involved in multi-modal ion transport are shown in the ribbon model (PDB code: 8JXO). Amino acids located in the intracellular region, extracellular region, and near the retinal are colored in orange, magenta, and yellow-green, respectively, in the stick representation. The retinal is shown in yellow. (**b**) Photocurrents of B1KCR variants in the presence and absence of metal cations. Representative photocurrent traces of B1KCR variants are shown. K–Na denotes recordings obtained with 110 mM K^+^ in the intracellular solution and 110 mM Na^+^ in the extracellular solution. NMG–NMG denotes recordings obtained with 110 mM NMG^+^ in both the intracellular and extracellular solutions. Blue bars indicate light illumination (480 nm, 6.4 mW mm^-2^). The membrane potential was stepped from −80 to +40 mV in 20-mV increments. Corresponding I–V relationships are shown. Black or gray indicates recordings obtained in the presence of metal cations, whereas blue indicates recordings obtained in their absence. (**c**) Key residues responsible for proton-pump function of B1KCR. The mechanism resembles very well those of BR.

An examination of K^+^ channel and H^+^ pump activities was conducted under the conditions of K^+^_in_/Na^+^_out_ and NMG^+^_in_/NMG^+^_out_, respectively (Extended Data Figs. 5 and 6). While studies have been reported on the K^+^ channel properties of other KCRs, such as HcKCR1 and WIChR1, no such reports are available for H^+^-pump properties. Among all the data, the results of the important mutants are displayed in Fig. 3b, whose photocurrent decays are shown in Extended Data Table 1. With respect to the K^+^ selectivity, the reversal potential was predominantly shifted by the mutations at the sites of the known K^+^ selectivity filter, including W101. The present study yielded further residues implicated in the K^+^ selectivity filter, including W83. With respect to the H^+^-pump activity, it was hypothesized that there would be an enhancement in activity for W101R. This is due to the fact that the corresponding residue of W101 in B1KCR is R82 in BR, a key residue in the H^+^ pump (Extended Data Fig. 2b)^5,53–55^. Although the H^+^ pump was retained in W83A, W101R exhibited an H^+^ channel rather than a pump (Fig. 3b).

D104 and D228 are located in proximity to the Schiff base in B1KCR (Extended Data Fig. 2b), whose mutations have been shown to impair both K^+^ channel and H^+^ pump activities (Fig. 3b). These results strongly suggest that D104 and D228 have a negative charge by deprotonation, as well as many microbial rhodopsins^5,6^. T108 is one of the DTD motif in B1KCR, and the T108A mutation significantly slowed down the kinetics, though K^+^ channel and H^+^ pump activities were maintained (Fig. 3b). A notable outcome was observed for the mutants of D115, another constituent of the DTD motif. As is known, D115N mutation largely alters the K^+^ selectivity, yet the sustained channel activity is observed. Conversely, the H^+^ pump activity was completely ablated (Fig. 3b). Therefore, D115 is a prerequisite for the H^+^ pump mode of B1KCR. D115 corresponds to D96 in BR, a H^+^ donor (Extended Data Fig. 1e)^5,56–58^, suggesting that D115 is a H^+^ donor in B1KCR. The activity of the H^+^ pump was rescued by the introduction of an aspartate near the position 115 (S65D-D115N). It can thus be concluded that the mechanism of the H^+^ pump in B1KCR bears a strong resemblance to those of the known outward H^+^ pumps, such as BR (Fig. 3c). Important role of the DTD motif was reported by means of laser flash-induced photocurrent experiments of cryptophyte channelrhodopsins^39,59,60^

We postulated two transport modes, K^+^ channel and H^+^ pump, to be isolated (Fig. 2a). However, if D115 donates a H^+^ to the Schiff base (Fig. 3c), H^+^ pump might occur even in the presence of K^+^. The validation of this possibility is best achieved through the implementation of ion-transport measurements, a process that utilizes a pH electrode^61^. In the course of the measurements, B1KCR-expressing yeast cells were exposed to light, and alteration in solvent pH were meticulously documented by means of pH electrode. In contrast to the control afforded by patch clamp measurements, the intracellular ion concentration remains unregulated (Fig. 4a). However, the transport of ions other than H^+^ can be measured by employing a protonophore, carbonyl cyanide 3-chlorophenylhydrazone (CCCP). In the context of a light-driven sodium pump KR2, sodium efflux is accompanied by a modest H^+^ movement in the opposite direction, which is augmented in the presence of CCCP (Extended Data Fig. 7a)^62^. A salient issue is that the incorporation of CCCP has the capability to either augment or diminish the observed pH fluctuations in the transportation of H^+^ or non-H^+^, correspondingly (Extended Data Fig. 7a). The experimental results demonstrated a substantial decrease in pH changes in the presence of 100 mM K^+^, Na^+^, and Mg^2+^ by the addition of CCCP (Fig. 4b, c and d), indicating the occurrence of an outward H^+^ pump.

**Fig. 4:**
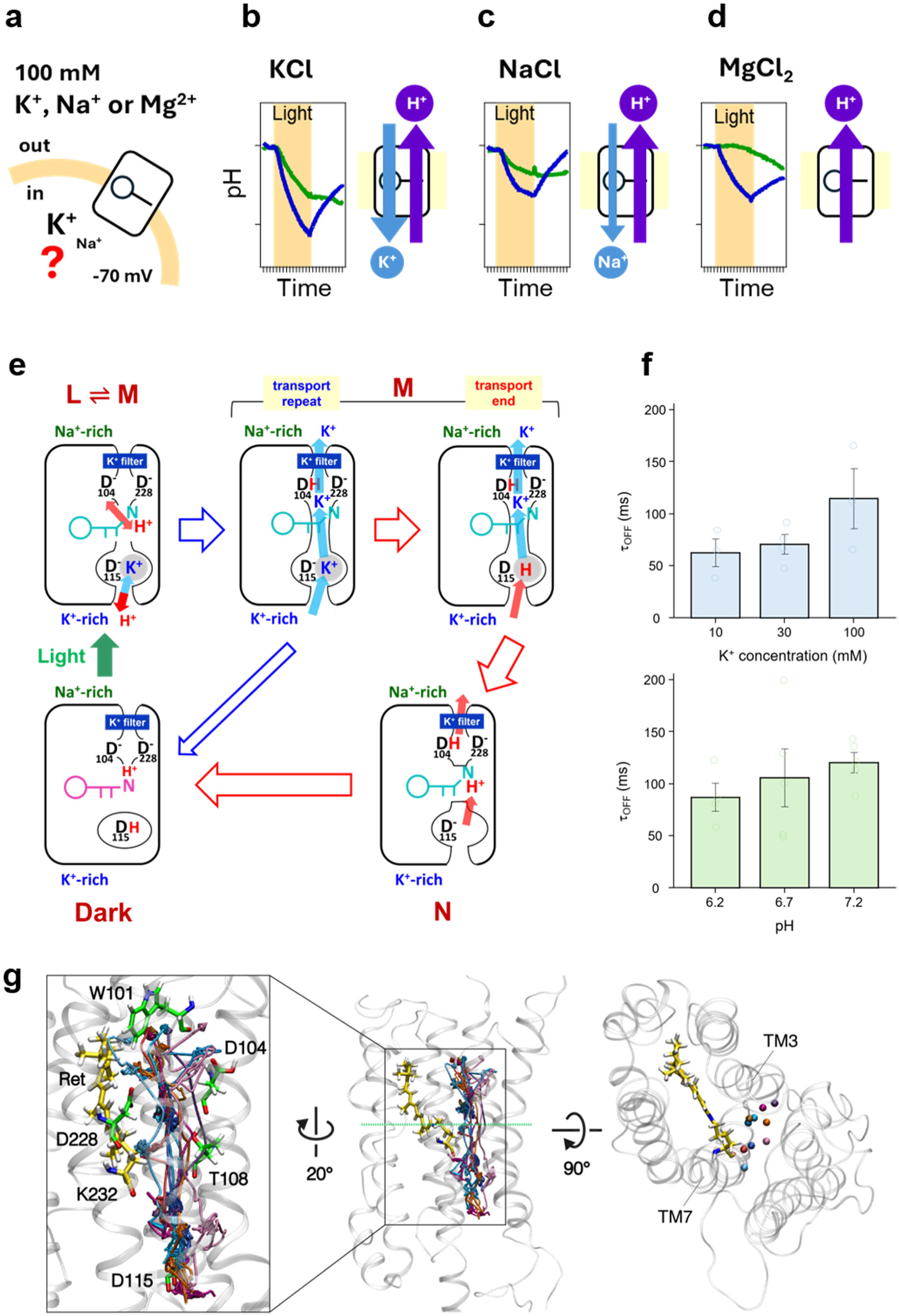
Multi-modal K^+^-channel and H^+^-pump functions in B1KCR. (**a**) Schematic illustration of the intracellular and extracellular ionic conditions of *P. pastoris* cells expressing B1KCR. (**b-d**) Light-induced pH profiles measured in the presence of extracellular 100 mM KCl (b), NaCl (c), or MgCl_2_ (d). Blue traces indicate the pH changes under each ionic condition, whereas green traces represent the corresponding measurements in the presence of CCCP. The right panels schematically illustrate the direction of ion transport under each condition. (**e**) Ion- transport model of B1KCR under physiological conditions. During the K^+^-channel function, proton uptake to D115 and the proton transfer to the Schiff base region cease the transport. (**f**) Time constants of photocurrent decay (τ_OFF_) of B1KCR measured under different intracellular K⁺ and H⁺ concentrations, with an extracellular solution containing 110 mM NaCl without K⁺ (pH 7.2). The cells were illuminated with light (λ = 470 nm). The holding potential was 0 mV. (**g**) K^+^ permeation pathways identified in MD simulations. The overall B1KCR structure is shown in side and top views, together with an enlarged view of the ion-conduction region. The spatial trajectories of the ten confirmed K^+^ permeation events, observed in six of the 30 independent 100-ns CompEL simulations (3.0 μs in total), are superimposed, each in a distinct color. The retinal chromophore and residues lining the K^+^ permeation route (W101, D104, T108, D115, D228, and K232) are shown as sticks.

The K^+^ transport mechanism in KCRs has been examined through the lens of a single sequential photocycle^36^. However, dual transports of K^+^ and H^+^ must be considered in the context of the B1KCR mechanism. Under the physiological conditions (K^+^_in_/Na^+^_out_), the intracellular K^+^ concentration is sufficiently high, and the incoming K^+^ probably passes near D115^36,37^, because the mutation influences the K^+^ selectivity. When the channel is open, K^+^ is transported through the pore, and a significant number of K^+^ are transported (transport repeat; Fig. 4e). However, if a H^+^ is bound to D115, and transferred to the Schiff base, such a H^+^ transfer completes the K^+^ transport (transport end; Fig. 4e). Consequently, the H^+^ pump mode functions as a blocker to the K^+^ transport function of KCRs (red arrow in Fig. 4e). The potential for spontaneous closure of K^+^ channels (blue arrow in Fig. 4e) is acknowledged, as previously documented.

The transport model depicted in Fig. 4E is predicated on the competitive binding of K^+^ or H^+^ to D115. In the event this is the case, the duration of transportation is contingent upon the concentrations of either K^+^ or H^+^. Accordingly, alterations were made to the intracellular ionic compositions, and the time constants of photocurrent decay (τ_OFF_) were measured. Consequently, τ_OFF_ exhibited an increase or decrease in duration in response to augmented K^+^ or H^+^ concentrations (reduction in pH) (Fig. 4f, Extended Data Fig. 7b, and Table 2). This observation lends support to our model, suggesting that the K^+^ transport may be influenced by intracellular concentrations of K^+^ and H^+^.

We then performed molecular dynamics (MD) simulations to examine a structurally feasible K^+^ permeation pathway. For this purpose, an open-state structure of B1KCR was built from that of HcKCR1 and K^+^ was initially bound near D115. By performing MD simulations comprising 30 independent 100-ns trajectories (3.0 μs in total) in the presence of transmembrane electrochemical gradient, ten complete K^+^ permeation events were identified during these simulations. Superposition of the trajectories revealed a continuous and broadly similar ion-conduction route through the protein, passing through the region lined by W101, D104, T108, D115, D228, and K232 (Fig. 4g). Analysis of the individual events showed that, although the spatial permeation routes were largely conserved, the time required for K^+^ to traverse the channel varied among events (Extended Data Fig. 8). This event-to-event variability is consistent with stochastic differences in the transient interactions of K^+^ with channel-lining residues during permeation. Representative time courses and the temporal distribution of all identified permeation events (Extended Data Fig. 8) support the structural feasibility of K^+^ permeation through B1KCR and suggest that variations in permeation time arise primarily from the dynamics of ion–protein interactions rather than from substantially different permeation routes.

## Light-dependent conversion from H^+^ pump to H^+^ channel

Microbial rhodopsins typically engage in photocycling reactions, while visual rhodopsins exhibit bleaching behavior in response to light absorption^5^. In this study, we identified a light-dependent conversion from H^+^ pump into H^+^ channel in B1KCR, which may be analogous to the bleaching of visual rhodopsins. In the patch-clamp recordings, rhodopsin-positive cells are typically identified by fluorescence from a fused fluorescent protein such as eYFP, whose illumination might influence ion transport, and this was indeed the case in the present study. Therefore, we endeavored to minimize exposure to fluorescence excitation light before and during the establishment of the whole-cell configuration (Extended Data Fig. 9a). The initial voltage-clamp recording of a predominantly dark-adapted B1KCR under NMG⁺ conditions yielded outward photocurrents, consistent with light-driven H⁺ pump (Fig. 5a). Two time constants were obtained for the photocurrent decay; 61.3 ms and 988 ms. However, upon repeated illumination of the same cell, the outward photocurrent exhibited a progressive decrease. Strikingly, from the fourth recording inward-directed photocurrents emerged at negative membrane potentials, whose decay was described by a single time constant (63.2 ms). This observation, clearly shown in the current–voltage (I–V) relationship (Fig. 5b), was unique under NMG⁺ conditions, as the K^+^-conducting state remains stable during repeated photoactivation (Fig. 5a and c).

**Fig. 5:**
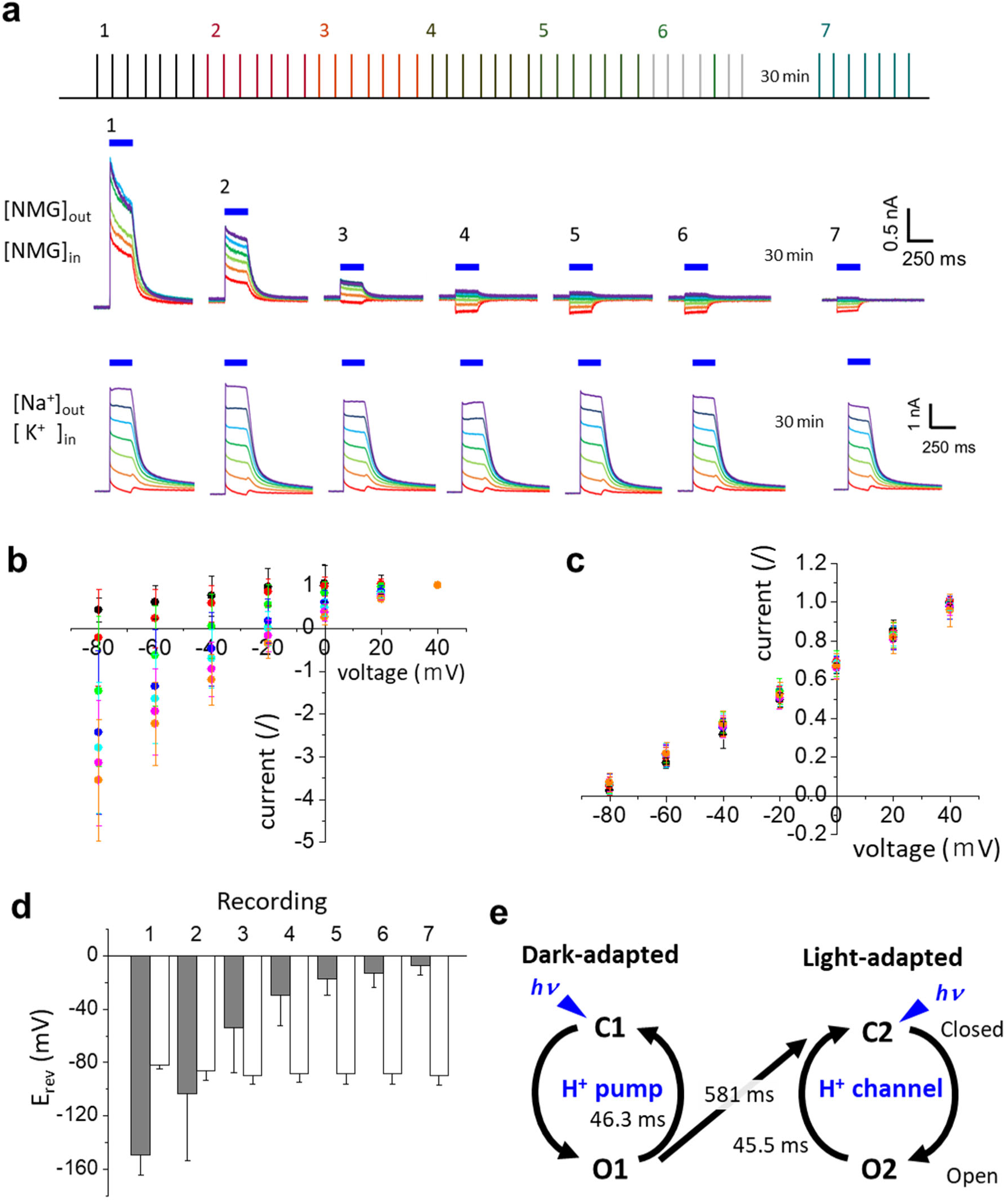
Light adaptation converts B1KCR from an H^+^-pumping mode to an H^+^-channel mode. (**a**) Repetitive photocurrent recordings from individual B1KCR-expressing cells under whole-cell voltage clamp. Photocurrent recordings were repeated six consecutive times in the same cell, followed by a 30-min dark interval and a seventh recording. Measurements were performed both in the absence and presence of metal cations. The comb-like schematic at the top indicates the timing of successive light-illumination protocols. Upper traces show consecutive recordings obtained in the absence of metal cations ([NMG^+^]_out_, [NMG^+^]_in_), whereas lower traces show those obtained in the presence of metal cations ([Na^+^]_out_, [K^+^]_in_). Blue bars indicate light illumination (480 nm, 6.4 mW mm^-2^). The membrane potential was stepped from −80 to +40 mV in 20-mV increments. (**b**) I–V relationships of B1KCR photocurrents recorded in the absence of metal cations ([NMG^+^]_out_, [NMG^+^]_in_). I–V relationships were constructed from each of the seven sequential recordings shown in A. For each I–V relationship, current amplitudes were normalized to the current measured at +40 mV, which was set to 1.0. (**c**) I–V relationships of B1KCR photocurrents recorded in the presence of metal cations ([Na^+^]_out_, [K^+^]_in_). I–V relationships were constructed from each of the seven sequential recordings shown in A. For each I–V relationship, current amplitudes were normalized to the current measured at +40 mV, which was set to 1.0. (**d**) Changes in the reversal potential during repeated photocurrent recordings from the first through seventh illumination protocol. Reversal potentials, defined as the membrane potential at which the photocurrent crossed zero, were determined from the seven I–V relationships shown in b and c. Gray, absence of metal cations ([NMG^+^]_out_, [NMG^+^]_in_); white, presence of metal cations ([Na^+^]_out_, [K^+^]_in_). (**e**) Schematic model for the functional transition of B1KCR from an H^+^-pump mode to an H^+^ - channel mode. In the dark-adapted state, B1KCR predominantly undergoes the C1–O1 photocycle and functions as an H^+^ pump. Repeated photoactivation progressively shifts the protein from the C1–O1 cycle toward the C2–O2 cycle, corresponding to the light-adapted state, in which B1KCR functions as an H^+^ channel. The time constants indicated in the scheme were derived from photocurrent recordings.

**Fig. 6:**
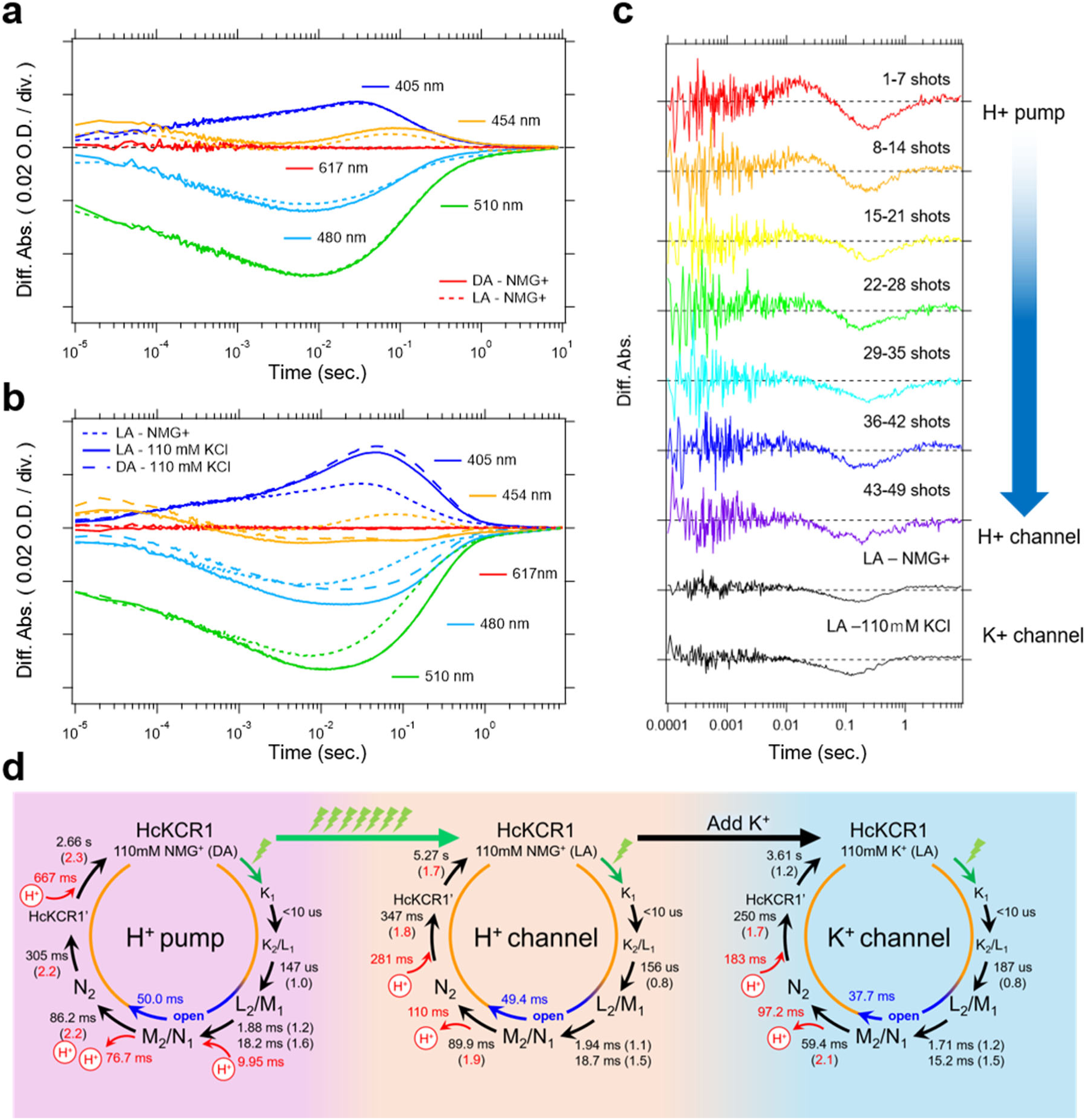
Photocycle kinetics and proton-transfer reactions of HcKCR1 revealed by flash photolysis. (**a**) Time-resolved absorbance changes of HcKCR1 monitored at 405, 454, 480, 510, and 617 nm in dark- and light-adapted H^+^-pump and H^+^-channel modes, respectively. The solid and dotted lines represent traces recorded in the dark- and light-adapted states. Trace colors correspond to the monitoring wavelengths. The absorbance scale of the y-axis is 0.02 O.D. per one division. The trace of the H^+^-pump mode was multiplied by 0.79 to normalize the reaction amplitude of the H^+^-channel mode. The x-axis is shown on a logarithmic scale. (**b**) Time- resolved absorbance changes of HcKCR1 monitored at 405, 454, 480, 510, and 617 nm in the light-adapted H^+^-channel mode, the dark-adapted K^+^-channel mode, and the light-adapted K^+^- channel mode. Dotted, dashed, and solid lines represent traces recorded in the light-adapted H^+^- channel, dark-adapted K^+^-channel, and light-adapted K^+^-channel modes, respectively. Trace colors correspond to the monitoring wavelengths. The absorbance scale of the y-axis is 0.02 O.D. per one division. The trace of the dark-adapted K^+^-channel mode was multiplied by 0.85 to normalize the reaction amplitude of the light-adapted K^+^-channel mode. The x-axis is shown on a logarithmic scale. (**c**) Time course of pyranine transient absorbance changes measured at 454 nm during the transition from the dark-adapted H^+^-pump mode to the light-adapted H^+^- channel mode, together with the response observed in the light-adapted K^+^-channel mode. Negative and positive signals indicate proton release from and proton uptake into HcKCR1, respectively. The absorbance scale is 0.005 O.D. per one division. (**d**) Photocycle schemes for each functional mode. Time constants were derived from the kinetic analyses shown in (a-c). Values in parentheses indicate the kinetic isotope effect (KIE). KIE values greater than 1.7 are highlighted in red, suggesting substantial proton-transfer involvement in the corresponding kinetic transitions. The D_2_O-based kinetic traces used for KIE determination are provided in the Supporting Information.

Under NMG⁺ conditions, the apparent reversal potential shifted progressively in the positive direction, from approximately −150 mV during the first recording to approximately −10 mV by the sixth recording (Fig. 5d). Whereas the strongly negative apparent reversal potential observed initially is characteristic of active outward H⁺ transport, the emergence of a reversal potential near the equilibrium potential expected for H^+^-conducting currents is consistent with passive H^+^ conduction. Thus, these observations indicate that repeated photoactivation converts B1KCR from an H⁺-pump mode to an H⁺-channel mode. It is important to note that umbllera- shaped I-V curves under NMG⁺ conditions, such as WiChR (Fig. 2c) and T108A B1KCR (Fig. 3c), may be attributed to the influence of the pump-to-channel conversion.

Based on these observations, we propose a reaction scheme for the H^+^-transport modes of B1KCR (Fig. 5e). In the dark-adapted state, B1KCR predominantly operates through a C1–O1 photocycle that exhibits active H⁺ pumping. Repeated photoactivation drives the protein into a light-adapted state associated with a distinct C2–O2 photocycle, in which B1KCR instead functions as H⁺ channel. It is noteworthy that the H⁺-channel persisted in the seventh recording performed after 30 min of dark incubation (Fig. 5a). Thus, recovery from the light-adapted C2 state to the original dark-adapted C1 state is extremely slow under our experimental conditions with no detectable functional recovery within 30 min. Photocurrent decays were similar between H⁺ pump and channel (∼60 ms), while we propose that the pump-to-channel transition occurs at ∼1 s, the slow component of photocurrent decay in the H⁺ pump (Fig. 5e).

This light-dependent conversion from H⁺ pumping to H⁺ channel was not exclusive to B1KCR. HcKCR1 exhibited a comparable, albeit accelerated, transition under NMG⁺ conditions (Extended Data Fig. 9b, d, e), indicating that this behavior is shared by distinct members of the KCR family. Furthermore, after HcKCR1 had entered the H⁺-channel mode in NMG⁺/NMG⁺ solutions, extracellular perfusion with a K⁺-containing solution converted the photocurrent being characteristic of K^+^-channel activity. These results demonstrate that the transport property of KCRs is determined by their photoadaptation state and the availability of permeant metal cations. This finding reveals an unusual functional plasticity in which the same rhodopsin can operate as an active H⁺ pump, an H⁺ channel, or a K⁺ channel depending on illumination history and ionic conditions. It is noteworthy that the slow component (988 ms in B1KCR) was not observed in the photocurrent decay of HcKCR1, which calls into question the conversion kinetics. The occurrence of a branching reaction with a time constant equal to that of the photocycle in HcKCR1 is a plausible hypothesis, which is supported by the accelerated transition from H⁺ pump to H⁺ channel in HcKCR1 (Extended Data Fig. 9b).

We subsequently proceeded to assess the multi-modality of KCRs in isolated proteins, which were expressed in yeast cells. Consequently, the obtained proteins were much smaller in amount for B1KCR than for HcKCR1 (Extended Data Fig. 10a). We thus applied flash photolysis to the purified proteins of HcKCR1. In this study, the samples were maintained in a state of darkness during preparation in either K^+^ or NMG⁺ solutions. The latter solution was exposed to illumination in NMG⁺ solution, followed by the subsequent addition of K⁺ solution. Identical absorption spectra were obtained for all four samples, dark-adapted (DA) NMG⁺ and K^+^ samples, and light-adapted (LA) NMG⁺ and K^+^ samples (Extended Data Fig. 10b). Time-resolved absorbance changes of HcKCR1 in NMG⁺ solutions were very similar between dark- and light-adapted samples (Fig. 6a and Extended Data Table 3), suggesting that distinction between the H^+^-pump and H^+^-channel modes was not clearly discernible through spectroscopic analysis. Conversely, photocycles exhibited variation between NMG⁺ and K^+^ samples, while uniform absorbance alterations were observed in the dark- and light-adapted K^+^ samples (Fig. 6b and Extended Data Table 4). Therefore, it can be concluded that the standard K^+^ channel mode of HcKCR1 is recovered by means of the addition of K^+^ into the H^+^-channel mode, a finding that was observed for B1KCR and HcKCR1 in patch clamp experiments. We obtained analogous results in D_2_O (Extended Data Fig. 10c), wherein the disparate photocycles observed in NMG⁺ and K^+^ samples were attributed to the presence of shared photointermediates, namely L, M , and N.

Although the photocycling dynamics exhibited identical behavior between the H^+^-pump and -channel modes of HcKCR1, flash photolysis measurements employing a pH sensitive dye pyranine yielded substantial results. Pyranine has a pKa at neutral pH, and pH changes in the solution can be monitored maximally at ∼450 nm. Transient absorbance changes at 454 nm were predominantly attributed to the photointermediates of HcKCR1 during repeated measurements. However, a discernible discrepancy was observed in the presence and absence of pyranine in solution (green and red lines, respectively, in Extended Data Fig. 10d). Double difference exhibited the positive and then negative kinetic feature for the dark-adapted NMG⁺ samples, while repeated illumination resulted in the only negative signal, which was also the case by the addition of K^+^ (Fig. 6c and Extended Data Table 5). These alterations in NMG⁺ solution are likely indicative of a transition from H^+^-pump to H^+^-channel mechanisms. The positive signal is indicative of elevated pH levels. The observed kinetic behavior of the H^+^ pump can be interpreted as follows; first an uptake of H^+^, followed by the release of two H^+^, and finally an uptake of a H^+^. Conversely, a H^+^ release and subsequent uptake occur in the H^+^ and K^+^ channels. In the NMG⁺ solution, the photocycle dynamics were found to be identical between the H^+^-pump and H^+^-channel modes, while the release and uptake of H^+^ were found to be different.

Present flash photolysis measurements successfully distinguished three functional modes of HcKCR1, H^+^ pump, H^+^ channel, and K^+^ channel (Fig. 6d). Photocycles were identical between H^+^ pump and channel, which was consistent with the same decay time of photocurrent in patch clamp experiment. Despite identical photocycle dynamics, additional H^+^ uptake and release were detected for H^+^ pump mode by the measurements using a pH-sensitive dye pyranine. The straightforward interpretation of the additional H^+^ uptake and release in the H^+^ pump is that they originate from the transported H^+^ in pump. On the other hand, there would be a group responsible for the H^+^ release and uptake in the H^+^ and K^+^ channels of HcKCR1, and such H^+^ circulation is possibly conserved in the H^+^ pump. From kinetic similarity with the H^+^ and K^+^ channels, we infer that the H^+^ movements in the H^+^ pump originate from (i) the circulation with release (76.7 ms) and uptake (667 ms), and (ii) the vectorial transport with the uptake (9.95 ms) and release (76.7 ms). In BR, the H^+^ release occurs before the uptake (Extended Data Fig. 1e), whereas the order is reversed (uptake is first) in many H^+^-pumping microbial rhodopsins because of the absence of the proton release group (PRG). The H^+^ pump in HcKCR1 appears to be also the case. Recent transient absorption study of HcKCR1 with pyranine exhibited the negative and then positive kinetic feature in K^+^ solution^36^, which differs from the present study (only negative feature) (Fig. 6c). The protein construct of HcKCR1 was common, while the expression systems were different; yeast and insect cells. Protonatable groups are possibly sensitive to microenvironment of HcKCR1.

## Discussion

This study reports on the multi-modal ion-transport function of KCRs, H^+^ pump and H^+^ channel, in addition to the previously identified K^+^ channel. Researchers concur that rhodopsins serve a fixed function during the course of evolution, but this is not the case. It was initially identified through electrophysiological experiments conducted on mammalian cells, and subsequently corroborated through spectroscopic analysis of purified HcKCR1 proteins following expression in yeast cells. Interestingly, the same photointermediates manifest with comparable kinetic components (Fig. 6), thereby implying that molecular events are likely to be analogous across the three functional states. Recent unitary conductance recordings of HcKCR1 and WiChR1 revealed the existence of multiple conductance states^52^, whose connection to the present observation is a subject worthy of further research. It is important to note that the functions of the H^+^ pump and H^+^ channel were originally identified in conditions that did not include K^+^, i.e. non-physiological conditions. However, detailed analysis has revealed that the H^+^ pump occurs even under physiological conditions, where cytoplasmic concentrations of K^+^ and H^+^ determine the closing time of the K^+^ channel. In this regard, the H^+^ pump in KCRs exerts a blocking effect on the conventional K^+^ channel. In the context of optogenetic applications of KCRs, the physiological H^+^ transport activity provides critical insights. The concurrent H^+^ flux has the capacity to induce local pH fluctuations and unexpectedly alter transmembrane electrochemical gradients. Consequently, it could influence the overall polarization state of target neurons alongside K^+^ conduction. Therefore, caution is warranted when deploying KCRs as optogenetic silencers to ensure that these unintended H^+^ dynamics do not confound the interpretation of physiological outcomes.

With regard to the H^+^ pump function of KCRs, the mutation study of B1KCR revealed a common mechanism to those of known H^+^ pumps, such as BR, including deprotonation at D104 and D228, and protonation at D115 (Fig. 3c). In this study, the H^+^ acceptor from the Schiff base was not identified, either D104 or D228, in B1KCR. Conversely, the H^+^ donor was identified as D115, due to the complete abolition of H^+^ pump by the D-to-N mutation, and the regained H^+^ pump activity by the additional S-to-D mutation near N115. T108 in the DTD motif of B1KCR exerts a significant influence on the kinetics of both the H^+^ pump and the K^+^ channel. This influence is attributed to the T-to-A mutation, which has been demonstrated to result in a substantial deceleration of the photocurrent decay. Consequently, the H^+^ pump function of KCRs is fully consistent with the conservation of the DTD motif. It is noteworthy that the H^+^- pump function was less evident in other channelrhodopsins than in KCR, suggesting the presence of other residues that are responsible for the pump function.

Irreversible conversion of H^+^ pump to H^+^ channel was an unexpected discovery in the present study, from which we proposed a parallel photocycle model with no reversion from H^+^ channel to H^+^ pump (Fig. 5). Parallel photocycle model has been established in the functional cycle of CrChR2, whose origin was the two configurations of the retinal chromophore, namely all-*trans* and 13-*cis*, 15-*syn* states^63^. Nevertheless, it seems improbable that the alteration between the H^+^ pump and channel originates from different chromophore structures, as the two states exhibit equivalent absorption spectra (Extended Data Fig. 9b), and chromophore structures of differing natures are ordinally in equilibrium by light. We infer that the conversion of the H^+^ pump to the H^+^ channel is accompanied by a modification of the specific protein structure, which is irreversible and cannot be reversed spontaneously to the original H^+^-pump structure. The observation of similar absorption spectra in the H^+^ pump and channel implicates that the portion under investigation is distant from the retinal chromophore. A hint was obtained from the mutation study of B1KCR. While a significant proportion of mutants exhibited H^+^- pump activity, W101R demonstrated H^+^ channel functionally even in the dark (Fig. 3b), thereby indicating that W101 plays a pivotal role in the conversion process. In the H^+^-pump mode, W101 probably changes the interaction for the uni-directionality, which is the case in R82 of BR, the corresponding residue. It is conceivable that W101 may be entrapped within another structure in the H^+^ channel.

The involvement of H^+^ pump during the K^+^ channel in KCRs was also an unexpected finding; however, this possibility was hypothesized on the basis that D115 plays an important role in the H^+^ pump and K^+^ channel of B1KCR. For this purpose, transport measurements employing a pH electrode represent a superior method, although this approach does not permit the control of intracellular ionic concentrations (Fig. 4a). The results unequivocally demonstrated the presence of H^+^ pump by B1KCR in K^+^ solution (Fig. 4b-d). Conversely, K^+^ channel activity exhibited in B1KCR was less pronounced in the pH electrode experiments. For instance, in the presence of extracellular Mg^2+^, only the outward H^+^ pump was measured, despite the presence of K^+^ inside yeast cells. The effective intracellular K^+^ concentration could be reduced in yeast cells under the present conditions.

The dual transports of K^+^ and H^+^ were modelled using of the competitive binding to D115, where the H^+^ binding functions as a blocker to the K^+^ transport function of KCRs (Fig. 4e). The model is supported by the increased or decreased τ_OFF_ in duration in response to augmented intracellular K^+^ or H^+^ concentrations (reduction in pH), respectively (Fig. 4f). It is reasonable to hypothesize that additional H^+^ uptake and release in the photocycle of H^+^ pump would be evident; yet the photocycle model of K^+^ channel does not consider the competitive binding of H^+^ (Fig. 6d). In order to facilitate a more profound comprehension of the subject, it is imperative to undertake a thorough and detailed analysis. One such technique is the patch clamp experiment, which involves the use of laser excitation. It is evident that laser patch clamp experiments are more effective in distinguishing between pumps and channels than continuous light illuminations, due to the fact that single photon reactions involve the transport of a single H^+^ in the pump, and multiple ions in the channels.

## Materials and Methods

### Genes

Rhodopsin genes of B1KCR from *Bilabrum sp.* (GenBank OP710242), KCRs from *Hyphochytrium catenoides*, HcKCR1 (MZ826862), HcKCR2 (MZ826861), HcCCR, WiChR from *Wobblia lunata* (OP710241), ChR2 from *Chlamydomonas reinhardtii*, GtACR1, GtCCR4^32^ and ChRmine^33^ were chemically synthesized (GenScript).

### Expression plasmids in mammalian cells

Plasmid DNAs of the channelrhodopsins, B1KCR, HcKCR1, HcKCR2, HcCCR, WiChR for expression in mammalian cells were created by an in-fusion reaction (In-Fusion® HD Cloning Kit, Takara Bio Inc., Shiga, Japan). pGtCCR4-3.0-eYFP was used as the template vector. The successful insertion of each ChR gene into the vector plasmid was verified by DNA sequencing (Fasmac Co., Ltd., Kanagawa, Japan).

### Cell culture

ND7/23 cells were grown on a coverslip in Dulbecco’s modified Eagle’s medium (Fujifilm Wako Pure Chemical Corporation, Osaka, Japan) supplemented with 2.5 μM all-trans-retinal and 5% fetal bovine serum under a 5% CO_2_ atmosphere at 37 °C. The expression plasmids were transiently transfected by using FuGENE HD (Promega, Madison, WI, USA) according to the manufacturer’s instructions. Electrophysiological recordings were then conducted 16–36 hours after transfection. Successfully transfected cells were identified by eYFP the fluorescence under a microscope prior to the measurements.

### Electrophysiology

Electrophysiological recordings were performed as described previously^64,65^. All experiments were carried out at room temperature (23 ± 2 °C). Photocurrents were recorded using an amplifier Axopatch 200B (Molecular Devices, San Jose, CA, USA) under a whole-cell voltage-clamp configuration. Data were filtered at 5 kHz, sampled at 10 kHz, digitized by Digidata1550 or 1550B (Molecular Devices), and stored in a computer. Irradiation at 470 nm or 530 nm was carried out using collimated LED (parts No. WLS-LED-0470-03-22 or LCS-0530-03-22, Mightex, Toronto, Canada) controlled by computer software (pCLAMP10.7, Molecular Devices). Light power was measured directly by an objective lens of an inverted microscope by a power meter or a visible light-sensing thermopile (LP1, Sanwa Electric Instruments Co., Ltd., Tokyo, Japan, or MIR-100Q, SSC Inc., Mie, Japan). The internal pipette solution in the absence of metal cation contained (in mM) 110 N-methyl D-glucamine (NMG), 2 MgCl_2_, 2 CaCl_2_, 10 HEPES, 10 EGTA, adjusted to pH 7.2 with HCl, while the one in the presence of metal cation contained (in mM) 110 K-gluconate, 1 NaCl, 2 MgCl_2_, 2 CaCl_2_, 10 HEPES, 10 EGTA, adjusted to pH 7.2 with KOH. The K⁺ concentration dependence was examined using the following solutions. (all concentrations are in mM): (1) 10 K-gluconate, 100 N-methyl D-glucamine, 2 MgCl_2_, 1 CaCl_2_, 10 HEPES, 10 EGTA, adjusted to pH 7.2; (2) 30 K-gluconate, 80 N-methyl D-glucamine, 2 MgCl_2_, 1 CaCl_2_, 10 HEPES, 10 EGTA, adjusted to pH 7.2; (3) 100 K-gluconate, 10 N-methyl D-glucamine, 2 MgCl_2_, 1 CaCl_2_, 10 HEPES, 10 EGTA, adjusted to pH 7.2. The extracellular solution contained (in mM) 110 N-methyl D-glucamine (NMG), 2 MgCl_2_, 2 CaCl_2_, 10 HEPES, adjusted to pH 7.2 with HCl, while the one in the presence of metal cation contained (in mM) 110 NaCl, 2 MgCl_2_, 2 CaCl_2_, 10 HEPES adjusted to pH 7.2 with NaOH. For measurements at different pHs, HEPES was replaced with MES (pH 6.0, 6.2 or 6.7). Time constants were determined by a single exponential fit unless noted otherwise.

Special care was taken to distinguish between the dark-adapted and light-adapted states of channelrhodopsins during photocurrent measurements and to minimize unintended light adaptation before recording (Fig. S9A). Following transfection, cells were maintained in darkness until electrophysiological measurements. During patch-clamp experiments, cells were observed using 600 nm transmitted light for routine visualization and for manipulation required to establish the whole-cell configuration. When fluorescence imaging was necessary to identify channelrhodopsin-expressing cells, excitation light was kept as weak as possible (69 µW mm^-2^) and applied only briefly, typically for 5–30 s, to minimize photoactivation before photocurrent recording.

## Statistical analysis

All data in the text and figures are expressed as the mean ± SEM.

### Expression plasmids in *Pichia pastoris*

The human codon-optimized genes encoding B1KCR and HcKCR1 were cloned into the pPICZB vector (Thermo Fisher Scientific, MD, USA) using the In-Fusion HD Cloning Kit (Takara Bio Inc., Shiga, Japan), resulting in proteins carrying a C-terminal His8-tag. Recombinant proteins were expressed in the *Pichia pastoris* strain SMD1168H (Thermo Fisher Scientific, MD, USA).

### Small-scale sample preparation and comparison of protein yields

Transformed strains were cultured in BMMY medium supplemented with 30 μM all-*trans* retinal for 24 h, and cells were harvested. The cells were disrupted using a fluidizer, and membrane fractions were prepared by ultracentrifugation. The membrane fractions were solubilized with 1% (w/v) DDM and 0.1% (w/v) CHS for 2 h, followed by ultracentrifugation to remove insoluble materials. The solubilized samples were purified by affinity chromatography using Co-TALON resin. Protein yields of B1KCR and HcKCR1 were compared by measuring their UV-visible absorption spectra.

### Ion-transport assay

Ion transport activity in *P. pastoris* cells was measured as previously described^61^. Cells were cultured in BMMY medium supplemented with 5% (w/v) DMSO and 30 μM all-*trans* retinal for 24 h. Cell density was estimated by measuring the optical density at 660 nm (OD660). A 7.5 mL cell suspension was adjusted to an OD660 of 4.0 and washed three times with 100 mM NaCl, 100 mM KCl, or 100 mM MgCl_2_. The washed cells were resuspended in the corresponding salt solution, kept in the dark, and illuminated with light at λ > 410 nm from an IMH-250A metal halide fiber illumination system (Sigma Koki, Japan) through a Y-43 glass filter (AGC Techno Glass, Japan). Light-induced pH changes were monitored using a pH electrode (HORIBA, Japan). Measurements were repeated under the same conditions in the presence of 30 μM carbonyl cyanide 3-chlorophenylhydrazone (CCCP), a protonophore.

### Sample purification from *Pichia pastoris* cells for flash photolysis

The transformed cells were precultured in BMGY medium for approximately 24 hours at 30°C. The pigment of HcKCR1 was expressed in the BMMY medium with a final concentration of 30 μM all-*trans-*retinal (Toronto Research Chemicals, Canada) for 24 hours at 30°C before harvesting. After induction, the expression medium was covered by foil, and isolation and preparation of membrane fractions were performed under dim red light without transferring the sample. The cells were collected at 4500 rpm, 10 min, 4°C condition and pellets were resuspended and treated with buffer A (50 mM HEPES, 150 mM NaCl, 150 mM KCl, 20 v/v% glycerol, pH 7.5) containing protease inhibitor (Sigma-Aldrich, Japan), wastase (TaKaRa, Japan), and DNase (Sigma-Aldrich, Japan) for >10 hours. The treated cells were disrupted by the 3-times passage through a high-pressure homogenizer (EmulsiFlex C3, Avestin, Inc., Canada). The disrupted cells were centrifuged at 35000 rpm, 10 min, 4°C in a fixed-angle rotor. The collected membrane pellets were resuspended in buffer A with 1.2% n-dodecyl-β-D-maltoside (DDM: anatrace) and stirred for >3 hours at 4°C. The solubilization mixture was centrifuged for 30 min at 100,000×*g* in a fixed-angle rotor at 4°C. The solubilized protein was applied to Co-TALON (QIAGEN, Germany), and the resin with bound the full-length HcKCR1 was washed with buffer B (50 mM HEPES, 150 mM NaCl, 150 mM KCl, 10 v/v% glycerol, pH 7.5) with 0.03% of DDM and then treated with buffer C (50 mM HEPES, 150 mM NaCl, 150 mM KCl, 300 mM Imidazole, 5 v/v% glycerol, pH 7.5). Purified samples were concentrated using a 50K MWCO concentrator and buffer exchanged into buffer D (110 mM NMG^+^, 2 mM CaCl_2_, 2 mM MgCl_2_, 10 mM EGTA, 10 mM HEPES-HCl, pH7.2) with 0.03% DDM, which is the same buffer condition as the patch clamp measurement. All purification procedures after solubilization are performed at 4°C or on ice in strictly controlled dim red light conditions. Purified samples are used for flash photolysis and transient H^+^ movement detection.

### Transient visible absorption measurements by nanosecond laser flash photolysis

For each transient absorption measurement, solubilized HcKCR1 samples were adjusted to an absorbance of 0.6 at the absorption maximum. For D_2_O measurements, samples initially prepared in H_2_O-based buffer D (110 mM NMG^+^, 2 mM CaCl_2_, 2 mM MgCl_2_, 10 mM EGTA, 10 mM HEPES-HCl, pH 7.2) containing 0.03% DDM were concentrated by ultrafiltration and subsequently diluted with D_2_O-based buffer D (110 mM NMG^+^, 2 mM CaCl_2_, 2 mM MgCl_2_, 10 mM EGTA, 10 mM HEPES-DCl, pD 7.2) containing 0.03% DDM. The pD in D_2_O buffer was calculated as apparent pH + 0.41. The final D₂O content of the samples was >95%. Dark-adapted data were obtained from samples that had been purified and stored under dark conditions for at least 24 hours. Light-adapted data were obtained from samples illuminated with continuous 530-nm light at an intensity of 0.17 mW mm^−2^ for 1 hour. For pyranine measurements, pyranine (Thermo Fisher Scientific Inc., USA) was dissolved in buffer D to prepare an 80 μM stock solution. The stock solution was added to protein samples to achieve a final pyranine concentration of 40 μM.

All sample preparation procedures were carefully performed in the dim red light condition. Each prepared sample was mounted on a quartz cuvette holder (Flash300, Quantum Northwest, USA). The measurement temperature was controlled at 20 °C using a thermostat bath (Alpha RA8, Lauda) with circulating water. Each sample was illuminated with a flash of second harmonics of a nanosecond-pulsed Nd^3+^−YAG laser (λ = 532 nm, INDI40, Spectra-Physics). The laser energy density was ∼2.5 mJ/cm^2^ per pulse, and the repetition rate was set to an ideal interval that was sufficiently slower than the photocycle rate of each sample to avoid unnecessary photoexcitation of the transient intermediates. The change in absorption after laser excitation was probed by monochromated light with wavelengths of 405, 454, 480, 510, and 617 nm from the output of an Xe arc lamp (L9289-01, Hamamatsu Photonics, Japan) through two monochromators (S-10, Soma Optics Ltd., Japan) placed in front and behind the sample cuvette. The change in intensity of the probe light that passed through the sample was monitored using a photomultiplier tube (R10699, Hamamatsu Photonics). For the pyranine measurement, 454 nm light was used as the probe light. The signal from the photomultiplier tube was averaged and stored using a digital-storage-oscilloscope (DPO7104 or MSO22 2-BW-500, Tektronix, Japan). Each trace was normalized to an identical reaction amplitude based on the absorption intensity observed before proton-transfer reactions. Each dataset of time-resolved transient absorption measurements was analyzed using Igor Pro ver. 9 and the Global Fit package with multiple exponential functions to obtain the decay time constants and pre-exponential factors.

## Structural analysis

The structural comparison and analysis were performed using Chimera X^66^.

## Phylogenetic analysis

Amino acid sequence alignments were generated by using MUSCLE^67^ and ClustalW^68^. Phylogenetic trees were inferred using the maximum-likelihood method implemented in IQ-TREE^69^, applying the LG + G + I substitution model in Figure 2B or the LG + G + I + F model in Figure S3A. Branch support values were estimated from 1000 ultrafast bootstrap iterations. Sequence data for the rhodopsins were obtained from the GenBank database.

## Molecular dynamics (MD) simulations

The initial open-state structure of B1KCR was constructed by homology modeling with MODELLER 10.8^70^, using the experimentally determined open-state structure (PDB ID: 9CDD^37^) as the template. Because the C-terminal region (residues 257 onward) was predicted to be unfolded in the homology model, these residues were removed from the final model used for MD simulations. Protonation states of titratable residues were assigned using PROPKA^71^. The retinal Schiff base was modeled in its deprotonated state, whereas Asp104 was modeled in its protonated form. Retinal was manually covalently linked to K232 via a Schiff-base.

The protein was described using the Amber ff14SB force field^72^. POPE lipids were modeled using the Lipid14 force field^73^, and the system was solvated with TIP3P water molecules^74^. Potassium, sodium, and chloride ions were described using the Joung–Cheatham monovalent ion parameters^75^ optimized for TIP3P water. Retinal and other non-standard molecules were parameterized using the general amber force field 2 (GAFF2)^76^.

To reproduce the experimentally observed ion-binding site, six K^+^ ions were initially placed within the protein. Their initial positions were determined by superimposing the open-state homology model onto the experimentally determined closed-state structure (PDB ID: 8JXO^40^) and placing the ions at the positions corresponding to the bound K^+^ ions observed in the experimental structure.

A double-membrane simulation system was constructed for CompEL calculations using GROMACS 2024.4^77^. The final simulation box measured approximately 100.3 × 100.6 × 269.2 Å^3^ and contained a total of 198,424 atoms. The membrane normal was aligned with the z-axis, and periodic boundary conditions were applied in all three dimensions.

Electrostatic interactions were evaluated using the particle mesh Ewald (PME) method^78^ with a real-space cutoff of 12 Å, while van der Waals interactions were truncated at 12 Å. Covalent bonds involving hydrogen atoms were constrained using the LINCS algorithm^79^, allowing an integration time step of 2 fs. The temperature was maintained at 300 K using the velocity-rescaling thermostat^80^, with separate coupling groups for the membrane/protein and solvent/ions. Pressure was controlled semi-isotropically at 1 atm using the C-rescale barostat^81^. Prior to production simulations, the systems were subjected to energy minimization followed by equilibration under the NVT and NPT ensembles.

A transmembrane electrochemical gradient was generated using the CompEL method^82^ implemented in GROMACS. The duplicated membrane system partitioned the simulation box into two aqueous compartments, A and B, and the desired ionic imbalance was maintained using the CompEL ion-swapping algorithm. Potassium and chloride ions were treated as exchangeable ion species. The target ion populations were set to 498 and 506 K^+^ ions in compartments A and B, respectively, and each compartment contained 492 Cl^−^ ions. Ion exchange was performed along the membrane normal (z-axis). Exchange attempts were made every 100 MD steps using a coupling interval of 10 steps and a threshold value of 1. A cylindrical exchange region with a radius of 6 Å and a height extending ±15 Å from the channel center was defined for each membrane copy, and only ions within these regions were eligible for exchange by the CompEL algorithm.

Production simulations were performed in the NPT ensemble at 300 K. Thirty independent 100-ns trajectories were generated using different initial velocity distributions, yielding a cumulative simulation time of 3.0 μs. Potassium permeation events were analyzed across all trajectories after removing periodic boundary artifacts and aligning the trajectories by least-squares fitting of the protein backbone atoms.

## Acknowledgments

We thank Y. Kanazawa (NIT) for assistance with electrophysiological experiments.

## Author contributions

S.P.T. and H.K. designed the research. R.S., M.K., S.T., J.T., K.M., S.H. and S.P.T. performed electrophysiological measurements using ND7/23 cells. R.A.Y. expressed B1KCR using *P. pastoris* cells, and measured ion-transport by pH electrode. T.N. analyzed structures and sequences with assistance from R.A.Y. and T.N. M.S. and R.A.Y. expressed KCRs using *P. pastoris* cells, and M.S. purified the proteins of HcKCR1. T.N., M.S. and Y.F. conducted flash photolysis. K.J.F. and T.Y. performed MD simulation. S.P.T., and H.K. wrote the manuscript with input from all authors. H.K. supervised the research.

## Competing interests

The authors declare that they have no conflicts of interest with the contents of this article.

**Correspondence and requests for materials**

should be addressed to Satoshi P. Tsunoda or Hideki Kandori.

## Data and materials availability

All other data are available within this article, its Supplementary Information and Source Data file, or from the corresponding author on reasonable request.

---

**Extended Data Fig. 1.**
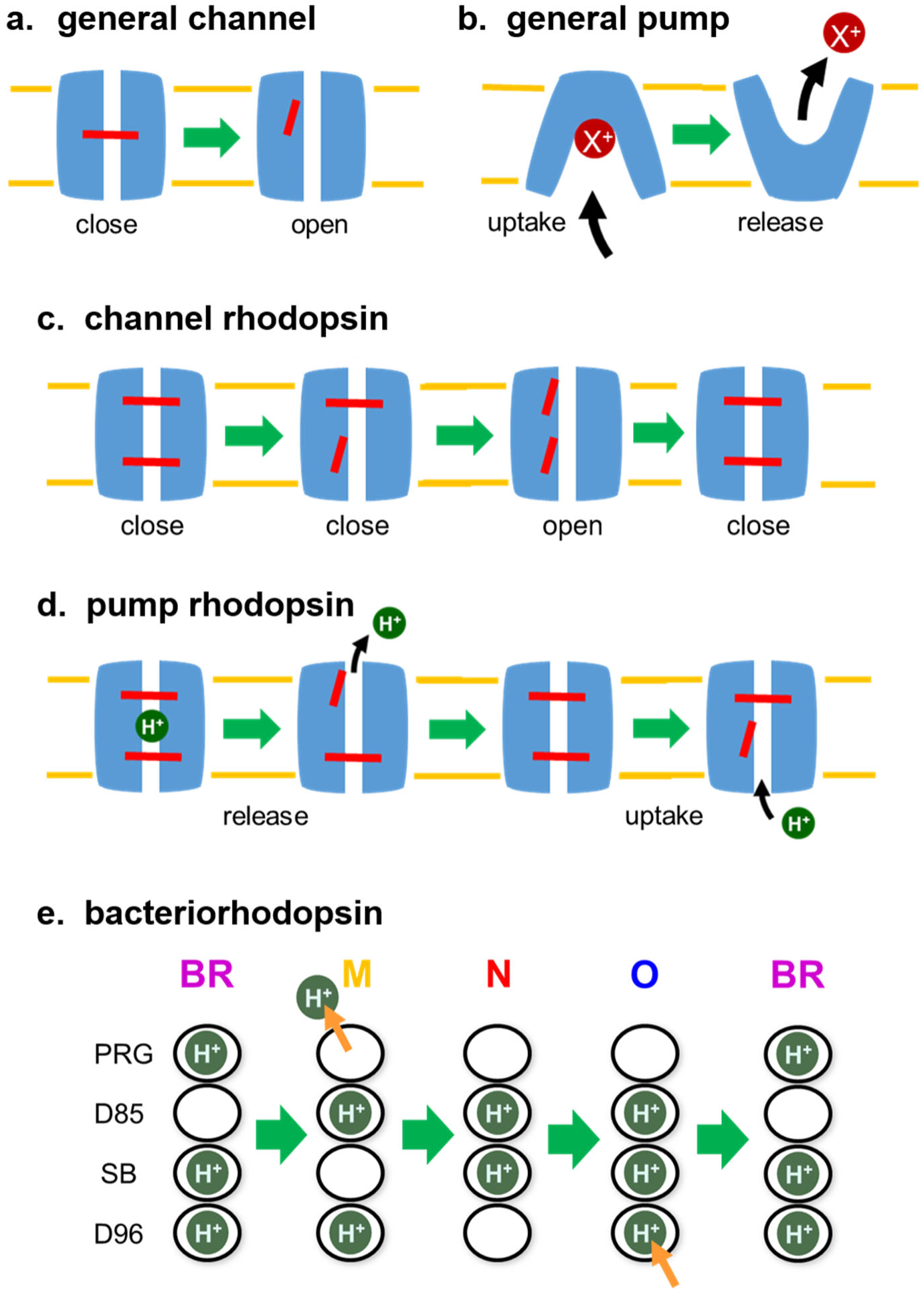
Mechanistic comparison of ion channel and pump. (**a**) General channel for passive ion transport. Ion channels are pore-forming membrane proteins, in which a gate controls open/close of channel. (**b**) General pump for active ion transport. Ion pumps perform vectorial transport of ions by use of energy, in which alternating access model explains the mechanism. (**c**) Passive ion transport mechanism of channel rhodopsin by use of two gates. If two gates are open at the same time, it becomes a channel. (**d**) Active ion transport mechanism of pump rhodopsin by use of two gates. If two gates are open in a specific order, it becomes a pump. (**e**) H^+^ pathway in a light-driven proton-pump bacteriorhodopsin (BR). SB and PRG stand for the Schiff base of the retinal chromophore and proton release group composed of E194, E204, and bound water molecules, respectively. In the dark state, protons are occupied by D96, SB, and PRG, and D85 is deprotonated. One proton is transferred as the results of five proton transfers inside BR during the photocycle containing M, N, and O intermediate.

**Extended Data Fig. 2.**
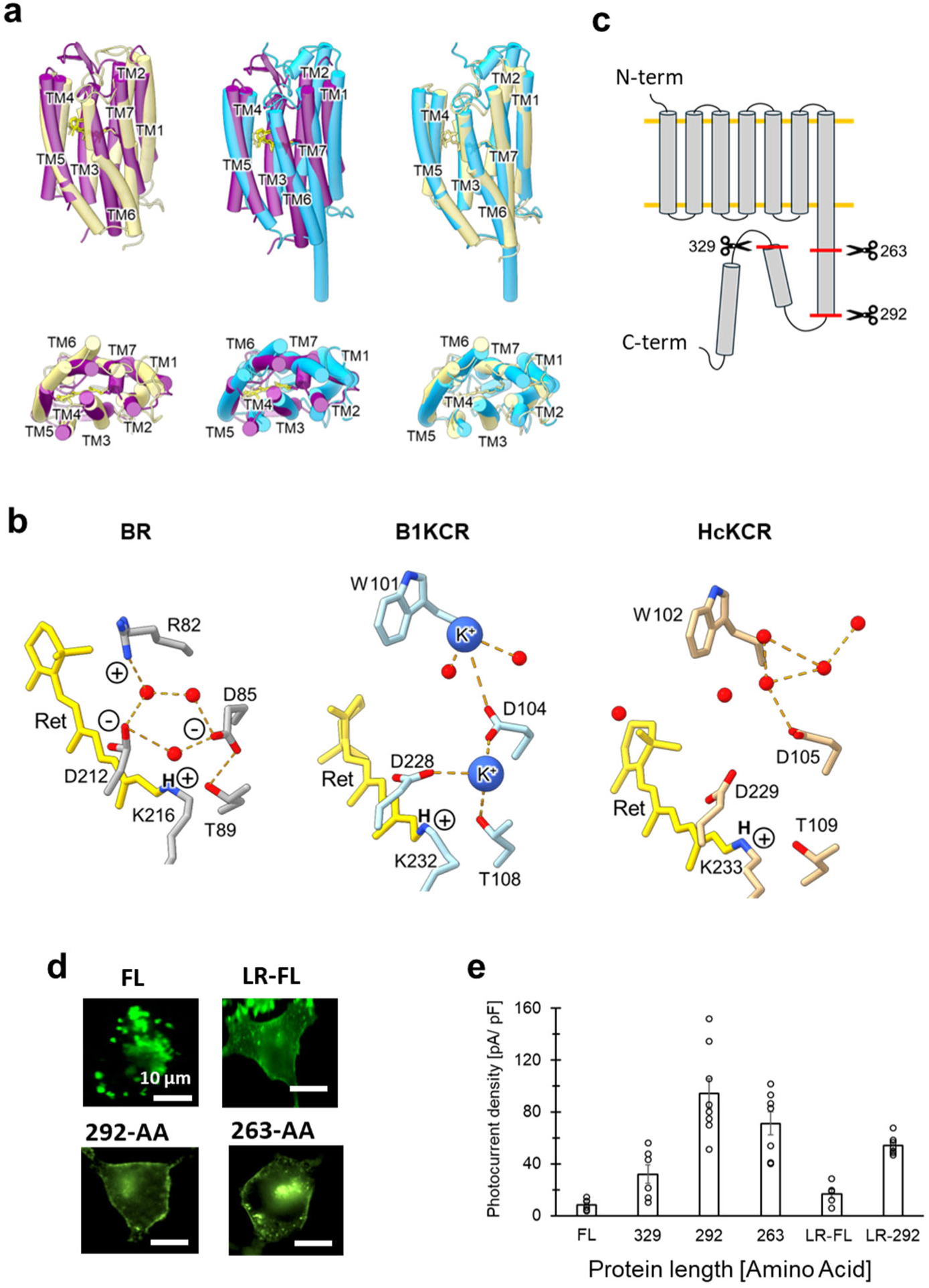
Structure and design of B1KCR. (**a**) Structural comparisons of BR (PDB code: 1M0M), B1KCR (8JXO), and HcKCR (9CDD), shown in dark violet, cyan, and khaki, respectively, in the cylinder representation. The top and bottom panels present the side view and the intracellular view, respectively. The retinal is shown in the stick representation. (**b**) Hydrogen-bond networks around the retinal, proton donor, and proton acceptor in BR, B1KCR, and HcKCR, displayed in the stick representation. Water molecules are shown as red spheres, and hydrogen bonds are indicated by orange dashed lines. Positively and negatively charged residues are highlighted. (**c**) Design of truncated B1KCR proteins at the C-terminal sides. (**d**) Cellular distributions of the fluorescence-tagged B1KCR proteins. (**e**) Photocurrent amplitudes of B1KCR constructs with different C-terminal lengths.

**Extended Data Fig. 3.**
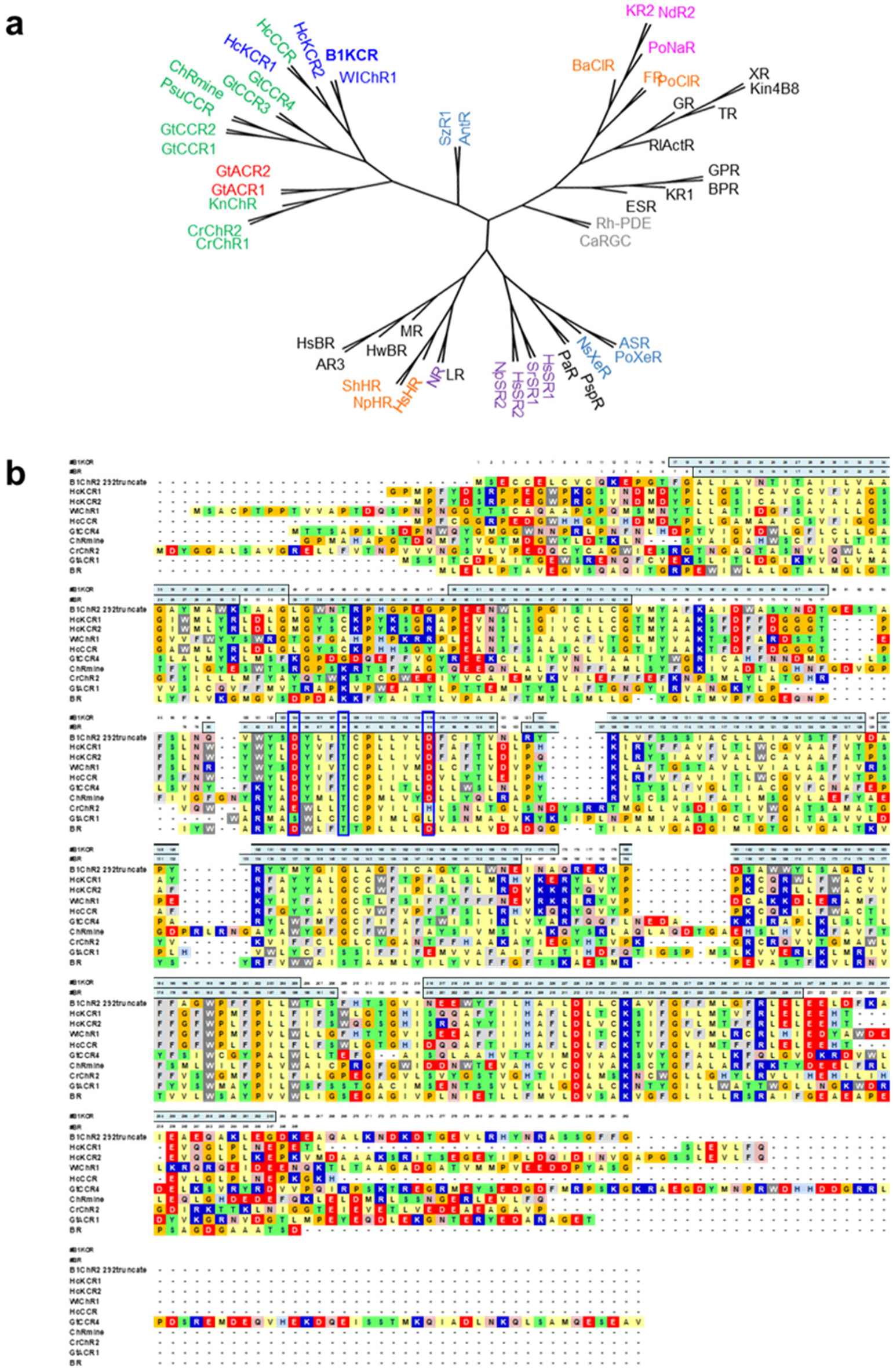
Phylogenetic tree and amino acid sequence alignment of the rhodopsins. (**a**) Phylogenetic tree of rhodopsins. Outward proton pumps, potassium channels, cation channels, anion channels, inward proton pumps, outward sodium pumps, inward chloride pumps, sensory rhodopsins, and enzyme rhodopsins are colored in black, blue, green, red, steelblue, pink, orange, violet, and gray, respectively. (**b**) Amino acid sequence alignment of the channel rhodopsins and BR.

**Extended Data Fig. 4.**
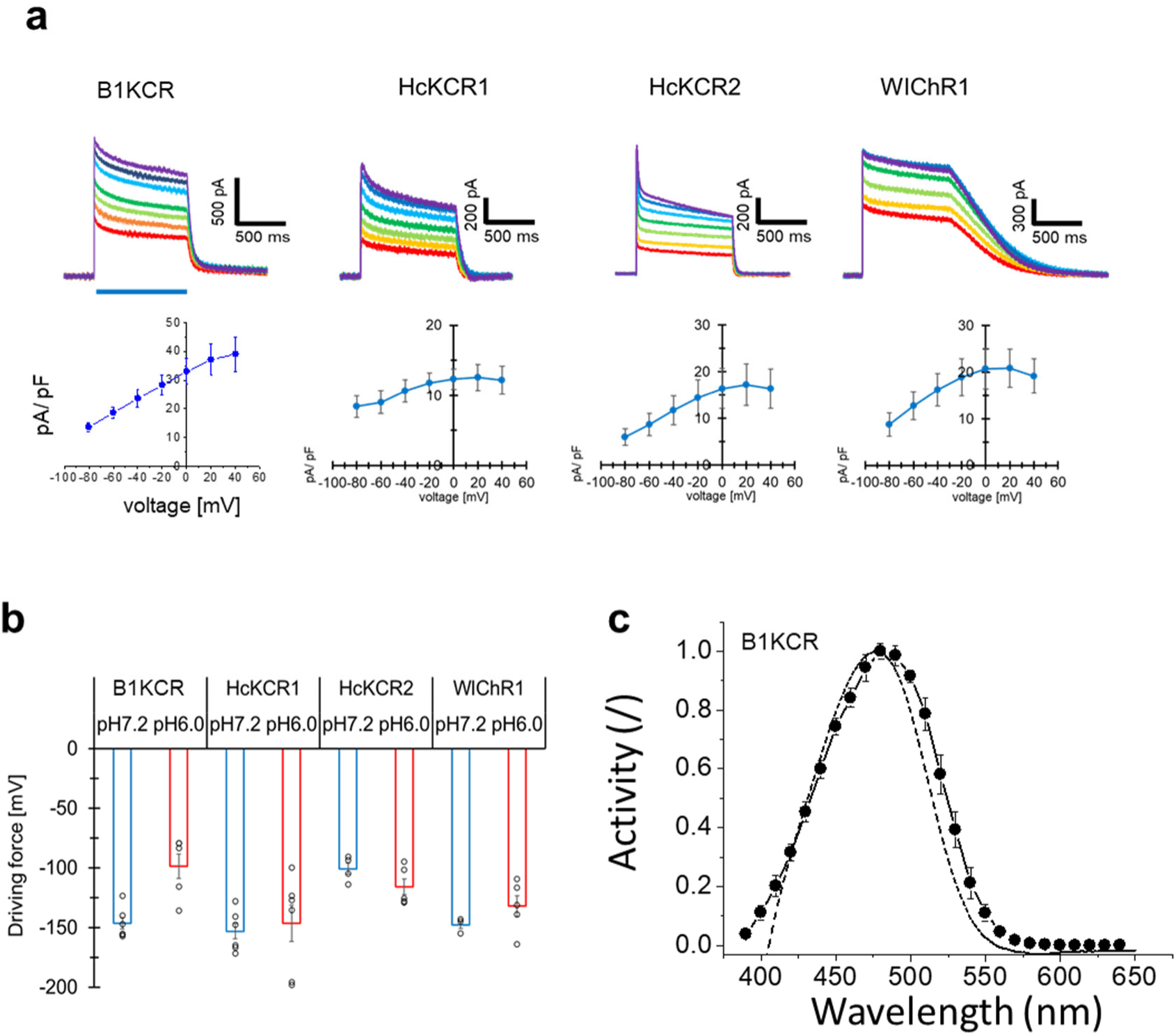
H^+^-transport of the KCRs. (**a**) Upper: Repetitive photocurrent recordings of the four KCRs in the absence of metal cation in solutions ([NMG^+^]_out_, [NMG^+^]_in_). The membrane potential was stepped from −80 to +40 mV in 20-mV increments. Blue bars indicate light illumination (480 nm, 6.9 mW mm^-2^). Lower: I–V relationships of the photocurrent. (**b**) Driving force of H^+^ transport in KCRs under a proton motive force. The driving force for proton pumping by each KCR was compared in the absence of an H+ concentration gradient (pH_in_ 7.2, pH_out_7.2) and in the presence of an H+ concentration gradient ((pH_in_ 7.2, pH_out_ 6.0). The driving force was defined as the membrane potential at which the photocurrent reached zero, estimated by linear extrapolation of the linear region of the photocurrent I–V relationship. (**c**) The action spectrum of B1KCR in the presence of metal cations in solutions [Na^+^]_out_, [K^+^]_in_. Photocurrent was measured by illuminating various wavelength of light. The membrane voltage was clamped at 0 mV. Dotted line: Absorption spectrum of the purified B1KCR.

**Extended Data Fig. 5.**
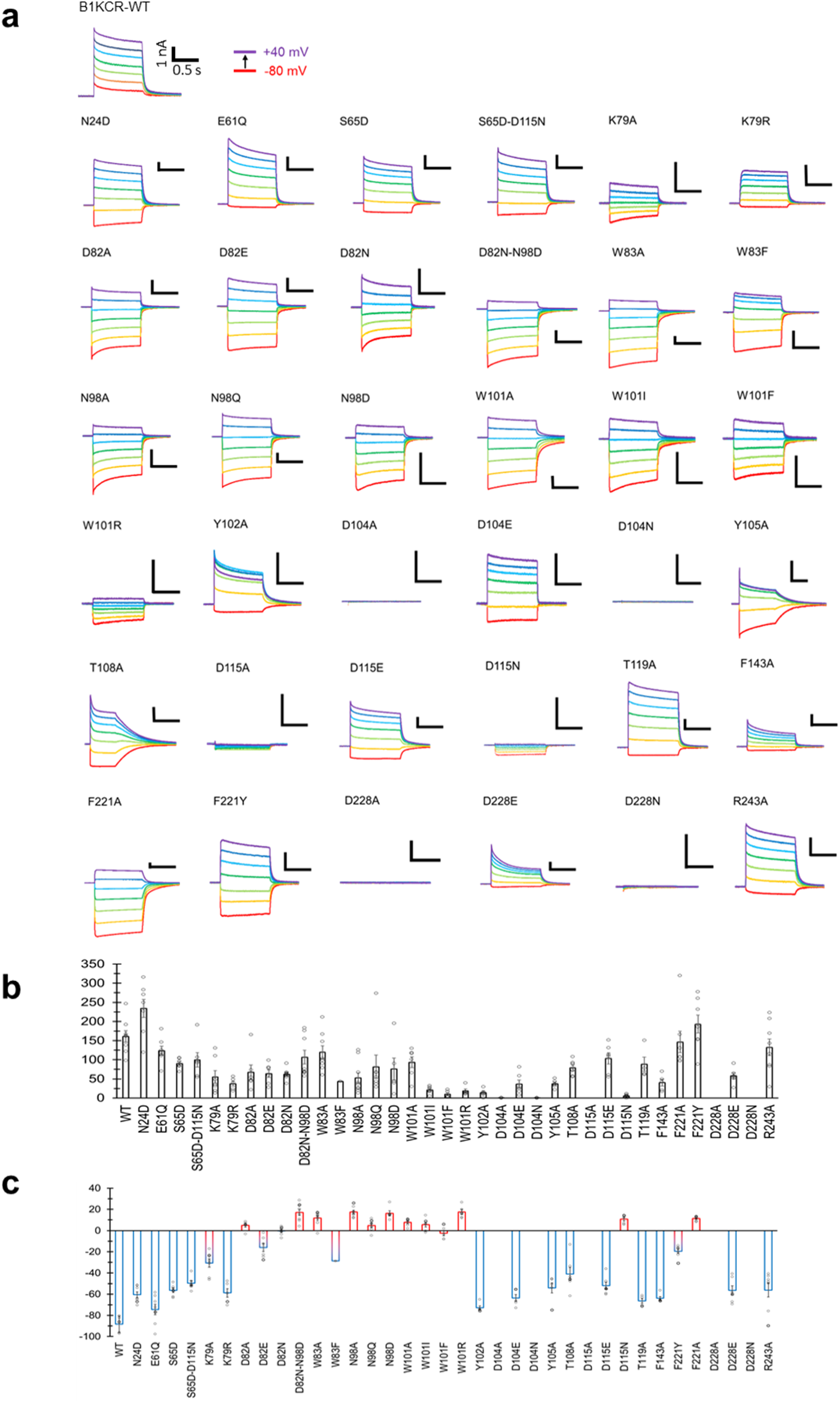
Photocurrent amplitudes and ion selectivity of B1KCR amino-acid substitution mutants. (**a**) Representative photocurrent traces of B1KCR mutants recorded with 110 mM K^+^ in the intracellular solution and 110 mM Na^+^ in the extracellular solution. Blue light (480 nm, 6.9 mW mm^-2^) was applied. The membrane potential was stepped from −80 to +40 mV in 20-mV increments. (**b**) Current densities (pA/pF) of B1KCR mutants. (**c**) Reversal potentials of B1KCR mutants.

**Extended Data Fig. 6.**
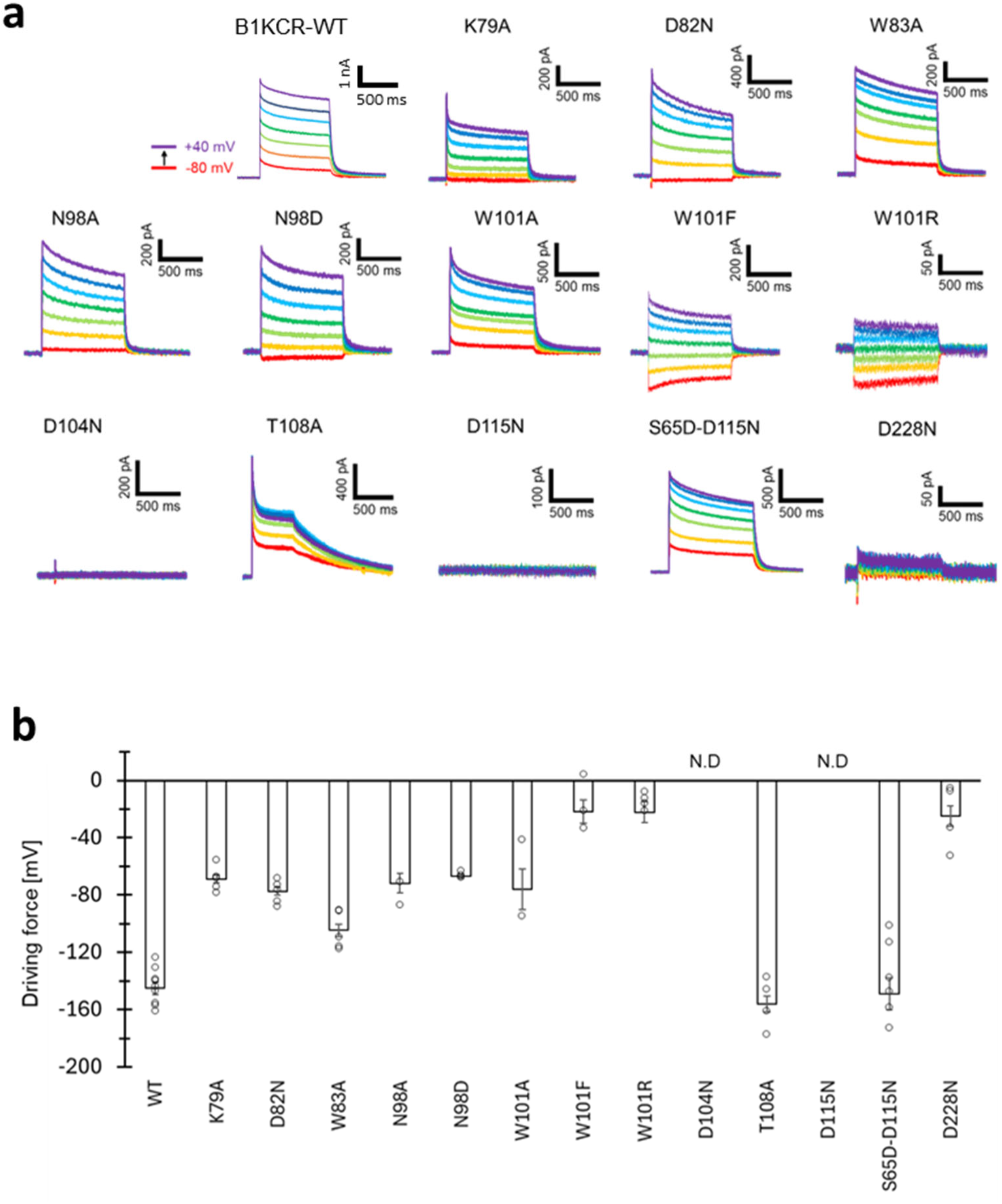
H^+^-transport properties of B1KCR amino-acid substitution mutants. (**a**) Representative photocurrent traces of the indicated B1KCR variants. Photocurrents of selected B1KCR mutants were measured in the absence of metal cations, with 110 mM NMG^+^ in both the intracellular and extracellular solutions, thereby isolating H^+^- dependent transport currents. No transmembrane pH gradient was imposed; both the intracellular and extracellular solutions were adjusted to pH 7.2. (**b**) Driving forces determined from the corresponding photocurrent measurements.

**Extended Data Fig. 7.**
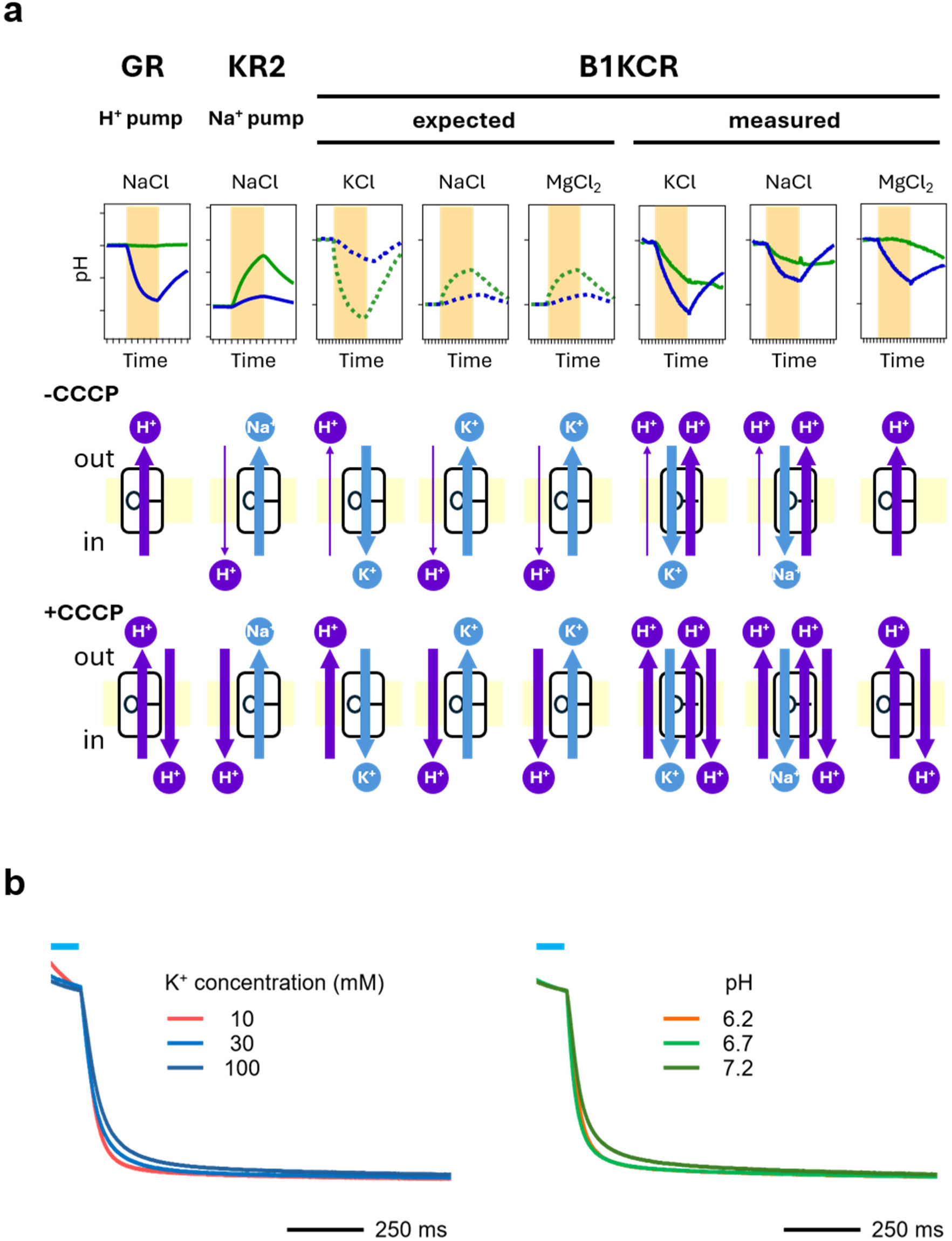
Ion transport activities of B1KCR under different ionic conditions. (**a**) Light-induced pH changes measured using *P. pastoris* cells suspended in the indicated salt solutions. Blue traces represent measurements in the absence of CCCP, whereas green traces represent the corresponding measurements in the presence of 30 μM CCCP. The middle and bottom panels schematically illustrate the proposed ion transport mechanisms inferred from the pH profiles in the absence and presence of CCCP, respectively. In the presence of 100 mM extracellular K^+^, it was hypothesized that there would be an influx of K^+^ by negative membrane potential, leading to a modest H^+^ movement outwardly (lowered pH) that would be augmented in the presence of CCCP (further lowered pH). The observation was precisely as anticipated in the absence of CCCP, whereas the results in its presence can be explained solely by the presence of an outward H^+^ pump. In the presence of 100 mM extracellular Na^+^ or Mg^2+^, it was hypothesized that there would be an efflux of K^+^ due to the low permeability to Na^+^ or the absence of permeability to Mg^2+^. This would result in a modest inward movement of H^+^ (increased pH). Nevertheless, a decrease in pH was observed, whose effect rendered less potent in the presence of CCCP. This necessitated the utilization of an outward H^+^ pump. In the case of Mg^2+^, the absence of transport with CCCP indicates the outward H^+^ pump exclusively. In the event of Na^+^, a decrease in pH with CCCP is indicative of non-H^+^ transport, which is most likely the result of an influx of Na^+^. (**b**) The decay phase of representative photocurrent traces of B1KCR under different intracellular K⁺ and H⁺ concentrations, with an extracellular solution containing 110 mM NaCl without K⁺ (pH 7.2). The cells were illuminated with light (λ = 470 nm) for the time, indicated by the blue bars. The holding potential was 0 mV.

**Extended Data Fig. 8.**
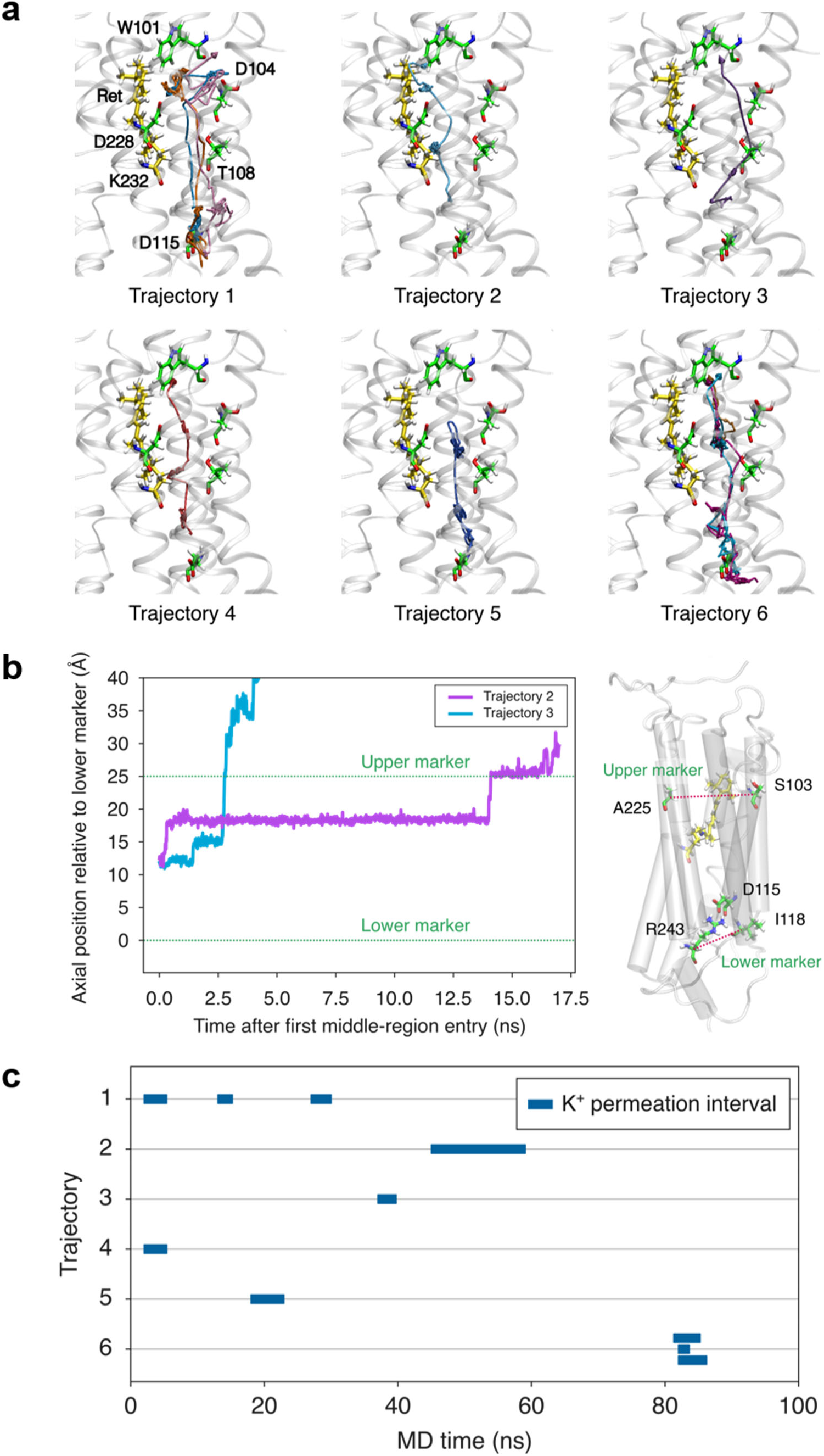
K^+^ permeation pathways and dynamics observed in MD simulations. (**a**) Spatial trajectories of the ten confirmed K^+^ permeation events observed in six of the 30 independent 100-ns simulations. The six subpanels correspond, from left to right and top to bottom, to trajectories 1–6. Multiple permeation events observed within the same simulation are superimposed in the corresponding subpanel, with each event shown in a distinct color. The protein is shown as a semitransparent white cartoon, the retinal chromophore (Ret) and K232 in yellow, and W101, D104, T108, D115, and D228 in green. The CompEL production simulations were initiated with K^+^ ions already present within the protein, with their positions differing among the equilibrated starting structures. Consequently, some permeation trajectories begin near D115, whereas others begin farther toward the retinal side and should therefore not be interpreted as complete K^+^ uptake pathways from the bulk solution. (**b**) Time evolution of the K^+^ position along the channel axis for two representative permeation events. Time zero is defined as the first entry of the K^+^ ion into the middle channel region. The axial position is measured from the lower channel marker, defined as the midpoint between the C_α_ atoms of I118 and R243, toward the upper channel marker, defined as the midpoint between the C_α_ atoms of S103 and A225; increasing values therefore correspond to the observed direction of K^+^ permeation. The structure on the right illustrates the positions of the upper and lower channel markers and D115. The two representative traces illustrate event-to-event variability in the time required for K^+^ to traverse the channel. (**c**) Timeline of the ten confirmed K^+^ permeation events identified across the 30 independent 100-ns trajectories. Each horizontal row corresponds to one of the six trajectories shown in (a), and each blue bar indicates the interval from the first entry of the K^+^ ion into the middle channel region to its first arrival in the upper region. Variation in bar length reflects differences in permeation time among individual events.

**Extended Data Fig. 9.**
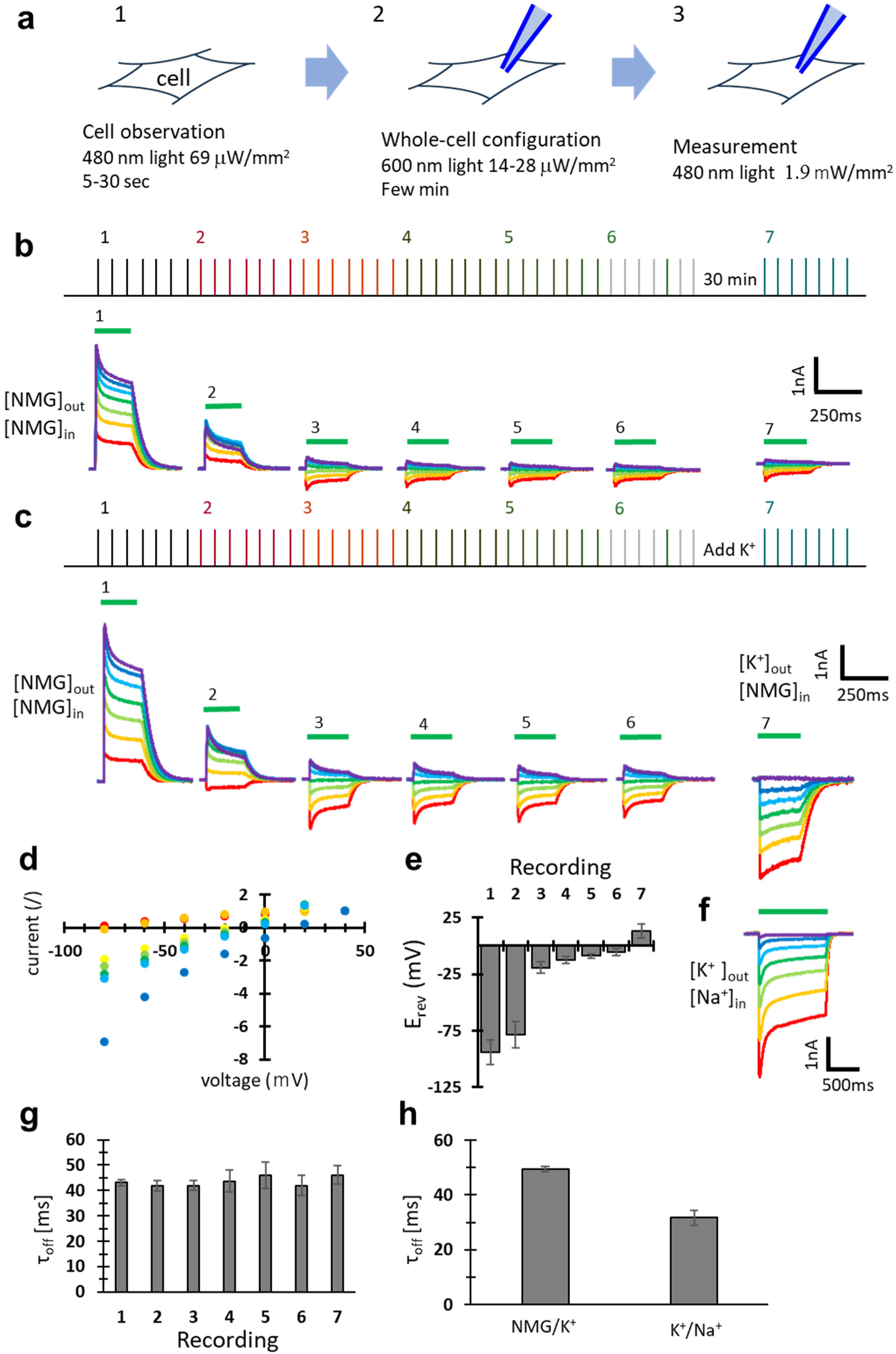
Light adaptation converts HcKCR1 from an H^+^-pumping mode to an H^+^-channel mode. (**a**) Precautions taken to minimize unintended light adaptation of channelrhodopsins during patch-clamp experiments. To avoid pre-adaptation, cells used for electrophysiological recordings were prepared under conditions that minimized light exposure. First, channelrhodopsin-expressing cells were identified by monitoring the fused fluorescent protein, using weak excitation light (480 nm, 69 µW mm^-2^) and a short observation period of 5 s to a maximum of 30 s. Second, cells were observed under 600-nm illumination during manipulation for establishment of the whole-cell configuration. Third, high-intensity 530-nm light (1.9 mW mm-^2^) was applied only at the onset of photocurrent measurements. (**b**) Repetitive photocurrent recordings from individual HcKCR1-expressing cells under whole-cell voltage clamp in the absence of metal cations ([NMG]out, [NMG]in). The membrane potential was stepped from −80 to +40 mV in 20-mV increments. Green bars indicate light illumination (530 nm, 1.9 mW mm^-2^). Photocurrent recordings were repeated six consecutive times in the same cell. The comb-like schematic at the top indicates the timing of successive light-illumination protocols. Traces show representative repetitive photocurrent recordings, followed by a 30-min dark interval and a seventh recording. (**c**) The same repetitive photocurrent recordings obtained under the same initial condition as B, followed by exchange of the extracellular solution to a K^+^-containing solution and subsequent measurement. (**d**) I–V relationships constructed from each of the seven sequential recordings shown in the photocurrent traces of b. For each I–V relationship, current amplitudes were normalized to the current measured at +40 mV, which was set to 1.0. (**e**) Changes in the reversal potential during repeated photocurrent recordings from the first through seventh illumination protocol. Reversal potentials, defined as the membrane potential at which the photocurrent crossed zero, were determined from the seven I–V relationships shown in d. (**f**) Photocurrent traces of HcKCR1 under the condition containing 110 mM K^+^ in the extracellular solution, and 110 mM Na^+^ in the intracellular solution. The membrane potential was stepped from −80 to +40 mV in 20-mV increments. Green bars indicate light illumination (530 nm, 1.9 mW mm^-2^). (**g**) Time constants of the photocurrent decay in the repetitive measurement in b. The data at 0 mV was analyzed. (**h**) Time constants of the photocurrent decay under two different solutions, shown in c (7^th^ measurement) and f, The data at 0 mV was analyzed.

**Extended Data Fig. 10.**
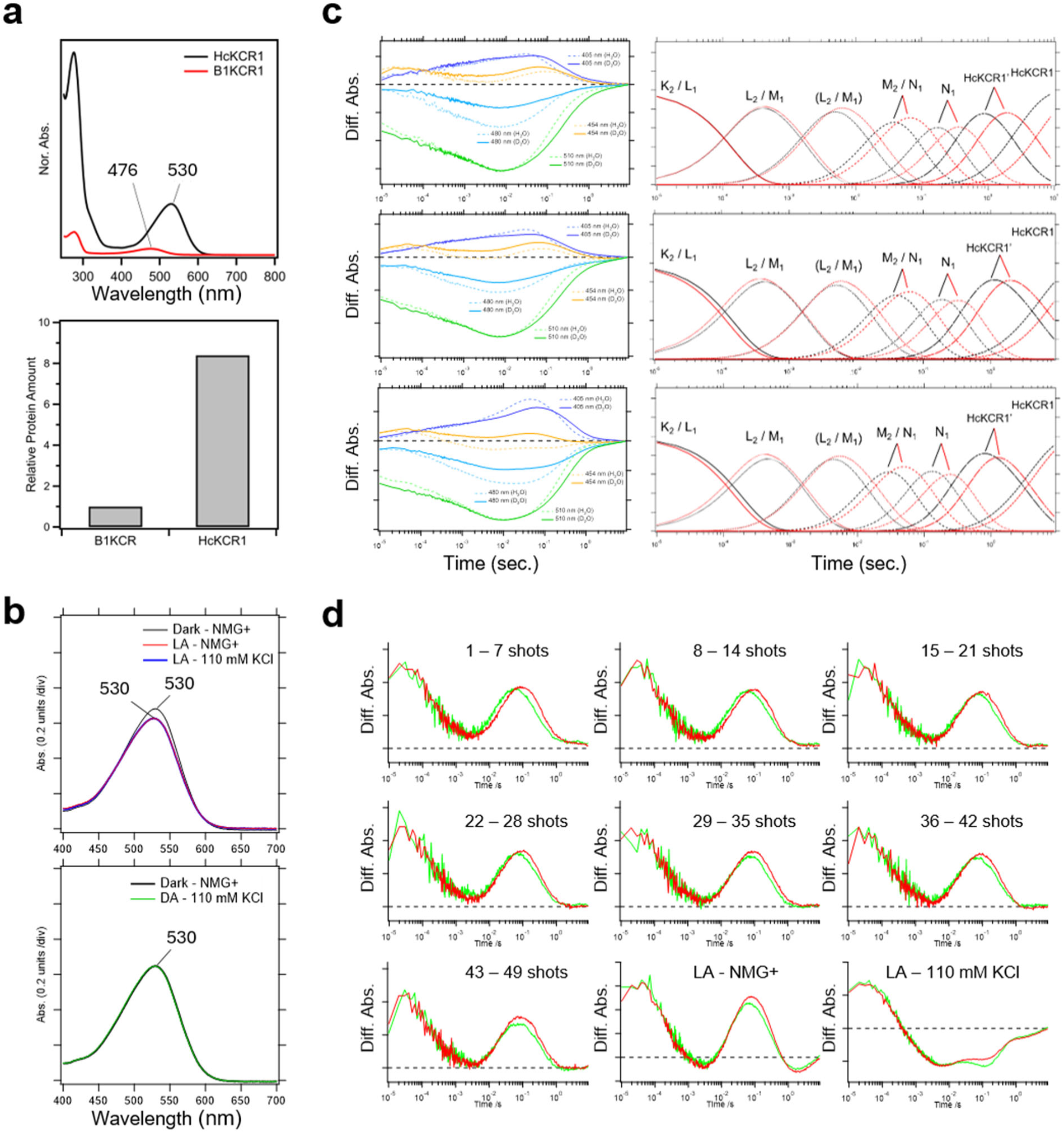
Purified KCR proteins and photoreaction dynamics of HcKCR1. (**a**) Comparison of the expression yields of B1KCR and HcKCR1. The upper panel shows the UV-visible absorption spectra of purified B1KCR and HcKCR1. The lower panel shows the relative protein yields estimated from the absorption spectra. (**b**; top panel) UV-vis spectra of the dark-adapted H^+^-pump (black line), the light-adapted H^+^-channel (red line), and the light-adapted K^+^-channel (blue line) modes. The λ_max_ of light-adapted modes are slightly blue-shifted, and their molar extinction coefficients are decreased. The light-adapted K^+^-channel (blue line) mode is multiplied by 0.98 to normalize the reaction amplitude of the light-adapted H^+^-channel mode. (**b**; bottom panel) UV-vis spectra of the dark-adapted H^+^-pump (black line) and the dark-adapted K^+^-channel (green line) modes. The dark-adapted K^+^-channel (green line) mode is multiplied by 0.95 to normalize the reaction amplitude of the light-adapted H^+^-channel mode. (**c, d, e**) (Left) Time-resolved absorbance changes of the dark-adapted H^+^-pump (c), the light-adapted H^+^-channel (d), the light-adapted K^+^-channel (e) modes monitored at 405, 454, 480, 510, and 617 nm in H_2_O and D_2_O buffer. Dotted and solid lines represent traces recorded in the H_2_O and D_2_O buffer, respectively. The absorbance scale of the y-axis is 0.02 O.D. per one division. The trace in the D_2_O buffer was multiplied by 0.73, 0.78, and 0.82 for (c), (d), and (e) to normalize the reaction amplitude in the H_2_O buffer, respectively. The x-axis is shown on a logarithmic scale. (Right) Kinetic isotope effects on the formation and decay of each intermediate in D_2_O buffer. (**f**) Raw data used for calculating pyranine transient absorbance changes. Each time trace was recorded in a buffer containing pyranine (green line) or in a buffer without pyranine (red line). Under the same conditions as patch-clamp recordings, seven laser- shot traces were averaged to generate a single trace during the transition from H^+^-pump mode to H^+^-channel mode. The light-adapted H^+^-channel and light-adapted K^+^-channel traces were generated by averaging 50 shots, respectively. The pyranine time courses in Fig. 6c were calculated from difference traces.

**Extended Data Table 1.**
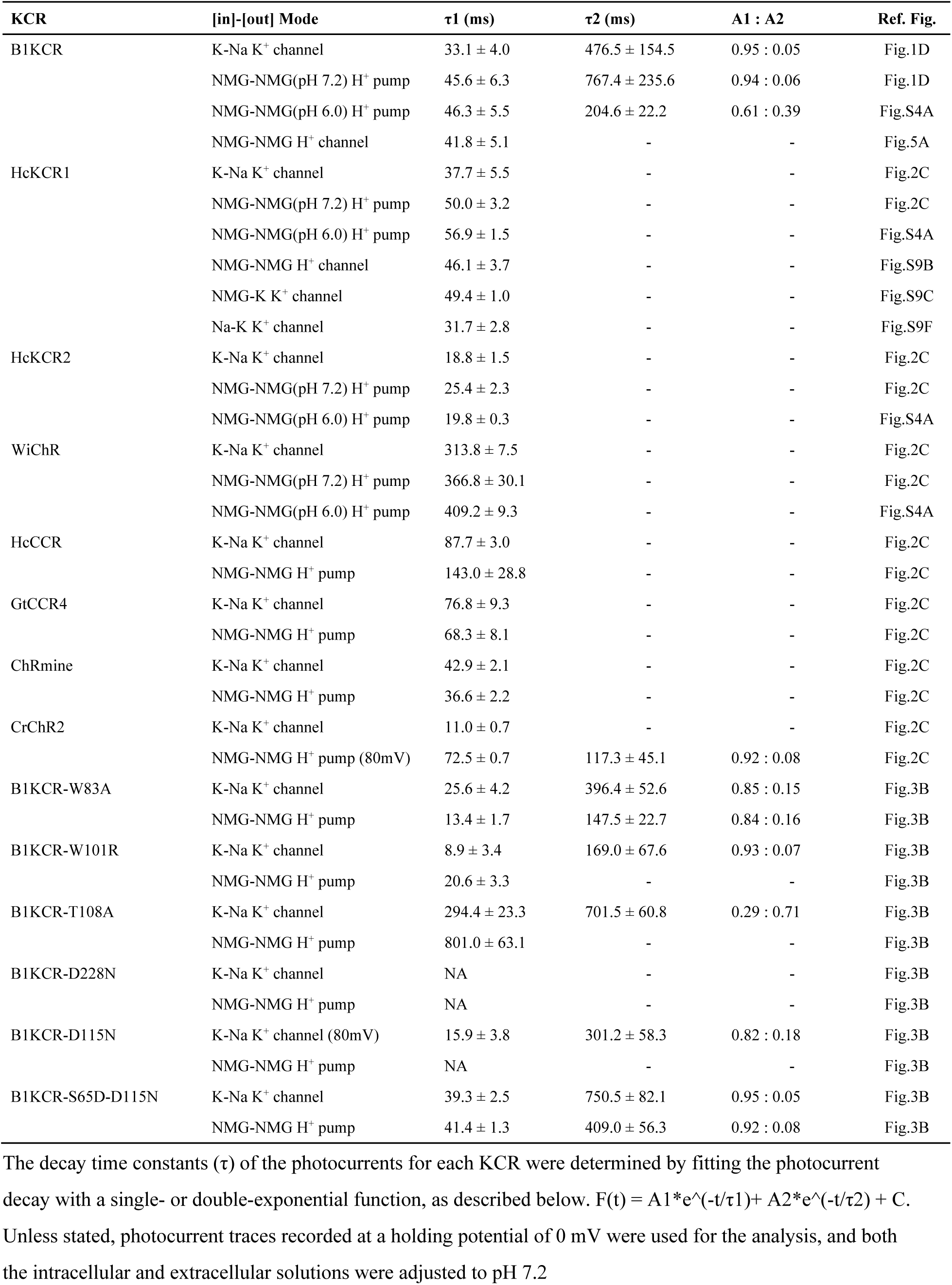
Time constants of KCR photocurrent decay determined by electrophysiological recordings

**Extended Data Table 2.** Time constants of B1KCR under cytoplasmic K^+^ and H^+^ concentrations obtained by electrophysiology

| Voltage (mV) | K<sup>+</sup> (mM) | pH | $\tau_1$ (ms) | $\tau_2$ (ms) | A1 : A2 | Ref. Fig. |
| --- | --- | --- | --- | --- | --- | --- |
| 0 | 10 | 7.2 | 35.0 ± 2.4 | 302.0 ± 64.4 | 0.9 : 0.1 | Fig.4F |
| 0 | 30 | 7.2 | 39.5 ± 2.85 | 297.4 ± 61.6 | 0.88 : 0.12 | Fig.4F |
| 0 | 100 | 7.2 | 42.4 ± 5.51 | 537.4 ± 131.2 | 0.86 : 0.14 | Fig.4F |
| 0 | 110 | 6.2 | 37.9 ± 1.6 | 591.9 ± 74.8 | 0.91 : 0.09 | Fig.4F |
| 0 | 110 | 6.7 | 36.3 ± 2.1 | 550 ± 86.2 | 0.88 : 0.12 | Fig.4F |
| 0 | 110 | 7.2 | 43.8 ± 2.3 | 475.6 ± 43.4 | 0.82 : 0.18 | Fig.4F |

The decay time constants ( $\tau$ ) of the photocurrents for each KCR were determined by fitting the photocurrent decay with a single- or double-exponential function, as described below. $F(t) = A1 * e^{-(t/\tau_1)} + A2 * e^{-(t/\tau_2)} + C$ .

**Extended Data Table 3.** Time constants and kinetic isotope effects from six-component global fitting in Fig. 6a and b

| Condition | Component | Assignment | $\tau_H$ (s) | $\tau_D$ (s) | KIE |
| --- | --- | --- | --- | --- | --- |
| DA - NMG+ | $\tau_1$ | K2/L1 → L2/M1 | 0.000147 ± 0.0000041 | 0.000145 ± 0.0000042 | 1.0 ± 0.040 |
| DA - NMG+ | $\tau_2$ | L2/M1 → M2/N1 (fast) | 0.00188 ± 0.000055 | 0.00218 ± 0.000042 | 1.2 ± 0.04 |
| DA - NMG+ | $\tau_3$ | L2/M1 → M2/N1 (slow) | 0.0182 ± 0.00061 | 0.0292 ± 0.00079 | 1.6 ± 0.1 |
| DA - NMG+ | $\tau_4$ | M2/N1 → N2 | 0.0862 ± 0.00240 | 0.186 ± 0.0067 | 2.2 ± 0.10 |
| DA - NMG+ | $\tau_5$ | N2 → HcKCR1' | 0.305 ± 0.016 | 0.671 ± 0.055 | 2.2 ± 0.2 |
| DA - NMG+ | $\tau_6$ | HcKCR1' → Dark | 2.66 ± 0.692 | 6.22 ± 1.797 | 2.3 ± 0.82 |
| LA - NMG+ | $\tau_1$ | K2/L1 → L2/M1 | 0.000156 ± 0.0000025 | 0.000126 ± 0.0000025 | 0.8 ± 0.020 |
| LA - NMG+ | $\tau_2$ | L2/M1 → M2/N1 (fast) | 0.00194 ± 0.000030 | 0.00208 ± 0.000027 | 1.1 ± 0.02 |
| LA - NMG+ | $\tau_3$ | L2/M1 → M2/N1 (slow) | 0.0187 ± 0.00043 | 0.0275 ± 0.00057 | 1.5 ± 0.04 |
| LA - NMG+ | $\tau_4$ | M2/N1 → N2 | 0.0899 ± 0.00139 | 0.167 ± 0.0042 | 1.9 ± 0.06 |
| LA - NMG+ | $\tau_5$ | N2 → HcKCR1' | 0.347 ± 0.0081 | 0.608 ± 0.027 | 1.8 ± 0.08 |
| LA - NMG+ | $\tau_6$ | HcKCR1' → Dark | 5.27 ± 0.49 | 8.81 ± 0.83 | 1.7 ± 0.22 |
| LA-KCl | $\tau_1$ | K2/L1 → L2/M1 | 0.000187 ± 0.0000028 | 0.000150 ± 0.0000027 | 0.8 ± 0.018 |
| LA-KCl | $\tau_2$ | L2/M1 → M2/N1 (fast) | 0.00171 ± 0.000022 | 0.00198 ± 0.000021 | 1.2 ± 0.02 |
| LA-KCl | $\tau_3$ | L2/M1 → M2/N1 (slow) | 0.0152 ± 0.0003 | 0.0228 ± 0.0004 | 1.5 ± 0.0 |
| LA-KCl | $\tau_4$ | M2/N1 → N2 | 0.0594 ± 0.00300 | 0.123 ± 0.0042 | 2.1 ± 0.12 |
| LA-KCl | $\tau_5$ | N2 → HcKCR1' | 0.250 ± 0.0013 | 0.421 ± 0.0044 | 1.7 ± 0.02 |
| LA-KCl | $\tau_6$ | HcKCR1' → Dark | 3.61 ± 0.225 | 4.42 ± 0.365 | 1.2 ± 0.12 |

Values are fitted values ± bootstrap SD. KIE was calculated as $\tau_D/\tau_H$ .

**Extended Data Table 4.** Time constants from six-component global fitting in Fig. 6b

| Component | Assignment | DA-KCl $\tau$ (s) | LA-KCl $\tau$ (s) |
| --- | --- | --- | --- |
| $\tau_1$ | K2/L1 → L2/M1 | $0.000179 \pm 0.000003$ | $0.000187 \pm 0.0000028$ |
| $\tau_2$ | L2/M1 → M2/N1 (fast) | $0.00174 \pm 0.000018$ | $0.00171 \pm 0.000022$ |
| $\tau_3$ | L2/M1 → M2/N1 (slow) | $0.0154 \pm 0.00024$ | $0.0152 \pm 0.0003$ |
| $\tau_4$ | M2/N1 → N2 | $0.0664 \pm 0.00234$ | $0.0594 \pm 0.00300$ |
| $\tau_5$ | N2 → HcKCR1' | $0.255 \pm 0.0013$ | $0.250 \pm 0.0013$ |
| $\tau_6$ | HcKCR1' → Dark | $2.30 \pm 0.199$ | $3.61 \pm 0.225$ |

Values are fitted values ± bootstrap SD. Time constants of DA-KCl and LA-KCl were determined in H<sub>2</sub>O buffer.

**Extended Data Table 5.** Pyranine time constants from exponential fitting in Fig. 6c

| Condition | Number of components | $\tau_1$ (s) | $\tau_2$ (s) | $\tau_3$ (s) |
| --- | --- | --- | --- | --- |
| DA | 3 | $0.00995 \pm 0.00214$ | $0.0767 \pm 0.0133$ | $0.667 \pm 0.116$ |
| LA | 2 | $0.110 \pm 0.0257$ | $0.281 \pm 0.0550$ | - |
| LA-KCl | 2 | $0.0972 \pm 0.0191$ | $0.183 \pm 0.0390$ | - |

